# *C. elegans* Sine oculis/SIX-type homeobox genes act as homeotic switches to define neuronal subtype identities

**DOI:** 10.1101/2022.04.19.488792

**Authors:** Cyril Cros, Oliver Hobert

## Abstract

The classification of neurons into distinct types reveals hierarchical taxonomic relationships that reflect the extent of similarity between neuronal cell types. At the base of such taxonomies are neuronal cells that are very similar to one another but differ in a small number of reproducible and select features. How are very similar members of a neuron class that share many features instructed to diversify into distinct subclasses? We show here that the six very similar members of the *C. elegans* IL2 sensory neuron class, which are all specified by a homeobox terminal selector, *unc-86/BRN3A/B*, differentiate into two subtly distinct subclasses, a dorsoventral subclass and a lateral subclass, by the toggle switch-like action of the SIX/Sine-oculis homeobox gene *unc-39. unc-39* is expressed only in the lateral IL2 neurons and loss of *unc-39* leads to a homeotic transformation of the lateral into the dorsoventral class; conversely, ectopic *unc-39* expression converts the dorsoventral subclass into the lateral subclass. Hence, a terminal selector homeobox gene controls both class-, as well as subclass-specific features, while a subordinate homeobox gene determines the ability of the class-specific homeobox gene to activate subtype-specific target genes. We find a similar regulatory mechanism to operate in a distinct class of six motor neurons. Our findings underscore the importance of homeobox genes in neuronal identity control and invite speculations about homeotic identity transformations as potential drivers of evolutionary novelty during cell type evolution in the brain.

**SIGNIFICANCE STATEMENT:** Anatomical and molecular studies have revealed that in many animal nervous systems, neuronal cell types can often be subclassified into highly related subtypes with only small phenotypic differences. We decipher here the regulatory logic of such cell type diversification processes. We show that identity features of neurons that are highly similar to one another are controlled by master regulatory transcription factors and that phenotypic differences between related cell types are controlled by downstream acting transcription factors that promote or antagonize the ability of such a master regulatory factor to control unique identity features. Our findings help explain how neuronal cell types diversify and suggest hypothetical scenarios for neuronal cell type evolution.

## INTRODUCTION

Understanding developmental programs in the nervous system necessitates the precise delineation and classification of neuronal cell types (Zeng and Sanes 2017). Single cell transcriptomic techniques have ushered in a new era of cell type classification in the nervous system of all animal species (Poulin et al. 2016; Network 2021). While classic anatomical and functional studies have revealed that neuronal cell types can be subdivided into distinct subtypes (Cajal 1911; Masland 2001), the inclusion of molecular criteria, and particularly recent scRNA transcriptome data, have begun to generate much finer-grained opportunities for cell type classification (Poulin et al. 2016; Zeng and Sanes 2017; Arendt et al. 2019; Network 2021). Such classification reveals taxonomic relationships with hierarchical structures that reflect the extent of similarity between neuronal cell types (Zeng and Sanes 2017; Arendt et al. 2019). For example, single cell profiling of retinal ganglion cells, which all share certain anatomical, functional and molecular features, identified novel subtypes and established hierarchical relationships between subtypes of previously known retinal ganglion cells (Shekhar et al. 2016). Similarly, recent comprehensive single cell transcriptomic analysis of other major parts of the brain (e.g. cortex and hippocampus) revealed a plethora of hierarchical, multi-tier subdivisions of specific inhibitory and excitatory neuron classes (Cembrowski and Spruston 2019; Network 2021; Yao et al. 2021). The hierarchical categorization of relationships of cellular identities poses an intriguing question, particularly at the bottom of such clustering diagrams: What are the molecular mechanisms by which cells that share a large swath of phenotypic identity features diversify into subtypes that express subtly distinct features? We address this problem here in the nervous system of the nematode *C. elegans*, where past anatomical (White et al. 1986) and recent scRNA-based molecular investigations (Taylor et al. 2021) have unearthed multiple example of sub-typologies of highly related neuronal cell types.

The six polymodal *C. elegans* IL2 sensory neurons, arranged as three symmetrically arranged neuron pairs (a dorsal, lateral and ventral pair; **Fig.1A**) have been classified together due to their characteristic cell body position, neurite projections, association with the inner labial sensilla and their shared synaptic connectivity (Ward et al. 1975; White et al. 1986)(**Fig.1B**). All six IL2 neurons stereotypically innervate the IL1 sensory-motor neurons and OLQ sensory neurons, the RIH interneuron and the RME and URA motorneurons, a pattern shared by no other neuron class (**Fig.1B**). However, only the lateral IL2 neuron pair, but not the dorsal or ventral pairs (from here on referred to as dorsoventral IL2 neurons) makes reciprocal chemical synapses with the dopaminergic ADE sensory neurons (**Fig.1B)**. In addition, the growth of extensive dendritic branches occurring during entry in a diapause arrest stage, when these neurons acquire specific mechanosensory functions (Lee et al. 2012), is mostly restricted to the dorsoventral IL2 neurons (Schroeder et al. 2013). Reporter genes studies as well as recent scRNA profiling have corroborated the notion that while all six IL2 neurons share the expression of a large number of genes, the dorsoventral IL2 and lateral IL2 neurons also differentially express a small number of genes (Wang et al. 2015; Hobert et al. 2016; Taylor et al. 2021)(**Fig.1C**). We investigate here how these subtype differentiation events are brought about. We show that the two IL2 subtypes represent two distinct, interconvertible states. Each state can be switched to the other by gain or loss of the SIX-type homeobox gene *unc-39*, which is normally expressed only in the lateral IL2 neurons; its removal leads to a homeotic identity transformation of IL2 lateral identity into dorsoventral IL2 identity, while ectopic expression of *unc-39* in the dorsoventral IL2 neurons switches them to the IL2 lateral identity. We find that another SIX homeobox gene, *ceh-32*, fulfills a similar role in another subtype diversification program of an unrelated, but also radially symmetric head motor neuron class.

**Fig 1:**
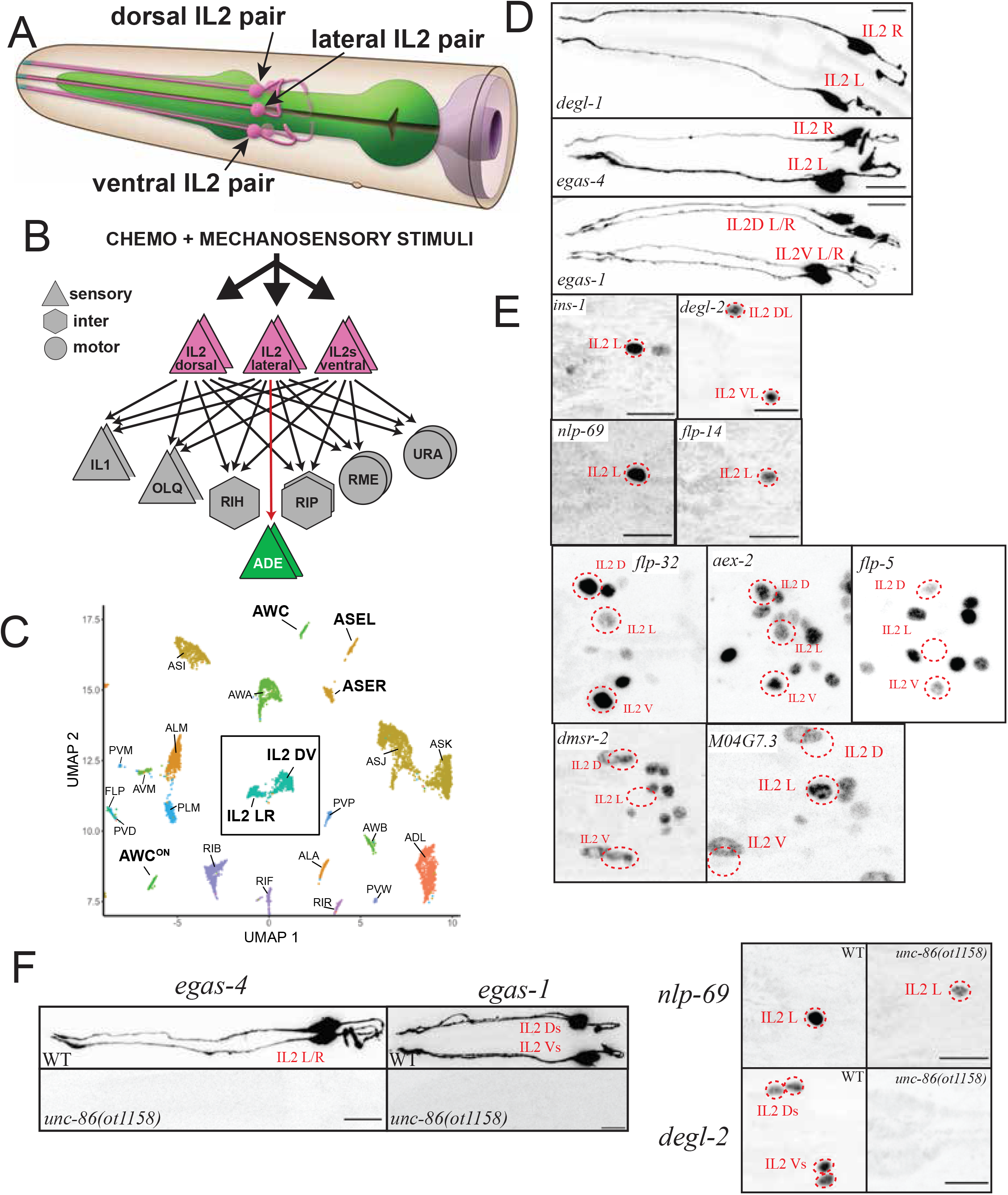
IL2 neurons are composed of distinct subclasses, all specified by the *unc-86* terminal selector. **A:** Schematic drawing of IL2 neurons, based on the original nervous system reconstruction (White et al. 1986) and reproduced with permission from Wormbook (Hall and Altun 2007). **B:** Synaptic connectivity of IL2 neurons. Data extracted from (White et al. 1986; Cook et al. 2019). **C:** Single cell transcriptome analysis showing similarity and differences among IL2 neurons subtypes, relative to other neuron classes. Reproduced from (Taylor et al. 2021). **D-E:** Expression pattern analysis using reporter alleles and/or reporter transgenes support IL2 subtype classification. **D:** Subtype-specific, cytoplasmically localized *gfp* promoter fusions expressed from integrated transgenes that show expression in the lateral IL2 subtype (*degl-1: otIs825, egas-4: otIs833)*, or the dorsoventral IL2 subtype (*egas-1: otIs846)*. **E:** Nuclear-localized reporter alleles for subtype-specific markers. Lateral IL2 subtype: *ins-1(syb5452[ins-1::SL2::gfp::H2B]), flp-14(syb3323[flp-14::T2A::3xNLS::gfp]), nlp-69(syb4512[nlp-69::SL2::gfp::H2B]), npr-37(syb4440[npr-37::SL2::gfp::H2B])*. Dorsoventral IL2 subtype: *degl-2(syb5229[degl-2::SL2::gfp::H2B]), flp-32(syb4374[flp-32::SL2::gfp::H2B]), aex-2(syb4447[aex-2::SL2::gfp::H2B]), flp-5(syb4513syb4513[flp-5::SL2::gfp::H2B]), dmsr-2(syb4514 [dmsr-2::SL2::gfp::H2B])*. For several of the more broadly expressed markers, NeuroPAL was used for cell identification (images in **Suppl. Fig.S2A**). **F:** IL2 subtype marker genes are *unc-86* dependent. In an *unc-86(ot1158)* null allele, both lateral IL2 markers (*nlp-69(syb4512), egas-4/otIs833)* and dorso ventral IL2 markers (*degl-2(syb5229), egas-1/otIs846, flp-32(syb4374), flp-5(syb4513))* are affected (Quantification in **Suppl. Fig.S2B**). Images and quantification for *flp-32* and *flp-5* regulation by *unc-86* are shown in **Suppl. Fig.S2C,D**.

## RESULTS

### Defining subtypes of the IL2 sensory neuron class

Single cell transcriptional profiling of the entire, mature nervous system of *C. elegans* (Taylor et al. 2021) revealed that all six IL2 neurons display a plethora of commonalities among the six neurons, supporting their initially anatomically-based classification into a unique class (White et al. 1986) (**Fig.1C**). Nevertheless, a distinctive set of molecules are differentially expressed between the dorsoventral pairs and the lateral pair (**Suppl. Fig.S1**)(Taylor et al. 2021). Since the IL2 neurons are possible mechanoreceptors, at least in the dauer stage (Lee et al. 2012), it is of particular note that scRNA profiling revealed distinctive expression patterns of the DEG/ENaC/ASIC family of ion channels, several of which (e.g., MEC-4, MEC-10) are known mechanoreceptors in other cellular contexts (**Suppl. Fig.S1**)(Al-Sheikh and Kang 2020). Specifically, scRNA data shows expression of two DEG/ENaC/ASIC family members, *asic-2* and *del-4*, in all six IL2 neurons, while five unusual DEG/ENaC/ASIC family members show a highly patterned expression in the IL2 subtypes (**Suppl. Fig.S1**). Three DEG/ENaC/ASIC proteins characterized by the existence of an additional EGF domain (the EGAS proteins)(Hobert 2013) are expressed either only in the dorsoventral IL2 neurons (*egas-1* and *egas-3*) or only in the lateral IL2 neurons (*egas-4*). Similarly, two small transmembrane proteins with an DEG/ENaC/ASIC domain also show expression in either the dorsoventral IL2 neurons (F58G6.8) or the lateral IL2s (Y57G11C.44)(Taylor et al. 2021). We named these two proteins DEGL-1 (Y57G11C.44) and DEGL-2 (F58G6.8). We generated promoter fusions for 3 of these 5 unusual, subtype-specific, putative mechanoreceptors (*egas-1, egas-4* and *degl-1*) and generated a reporter allele by CRISPR/Cas9 genome engineering for another one (*degl-2)*. These reporters confirmed their expression in either the lateral or dorsoventral IL2 subtypes, as predicted by the scRNA analysis (**Fig.1D, E**).

In addition, as per our scRNA data (Taylor et al. 2021), several neuropeptides and neuropeptide receptors also show IL2 subtype-specific expression (**Suppl. Fig.S1**). We recapitulate the scRNA data by generating reporter alleles (via CRISPR/Cas9 genome engineering) for four neuropeptides (*flp-32, flp-5, flp-14, nlp-69*), two neuropeptide receptor (*dmsr-2, M04G7*.*3/npr-37*) and one insulin-like peptide (*ins-1)*. We found that these genes are indeed all expressed in a subtype specific manner in either lateral (*flp-14, nlp-69, ins-1, M04G7*.*3/npr-37*) or dorsoventral IL2 neurons (*flp-32, flp-5, dmsr-2, aex-2*)(**Fig.1D,E, Suppl. Fig. S2A**). Taken together, the dorsal and ventral pair form one subclass of IL2 neurons and the lateral pair another subclass.

### Subclass specific features are controlled by the class-specific terminal selector *unc-86*

The identity of all six IL2 neurons is specified by the Brn3A-type POU homeobox gene *unc-86*, a phylogenetically conserved, terminal selector transcription factor, called BRN3 in vertebrates (Shaham and Bargmann 2002; Zhang et al. 2014; Leyva-Diaz et al. 2020). UNC-86/BRN3 is expressed in all six IL2 neurons and is required for the expression of all tested molecular features of the IL2 neurons (Finney and Ruvkun 1990; Shaham and Bargmann 2002; Zhang et al. 2014), as well as for their unique dendritic morphology (Schroeder et al.2013). The specificity of action of UNC-86/BRN3, which is expressed in multiple other neuron types as well (Finney and Ruvkun 1990), is determined by the IL2-neuron-specific collaboration of UNC-86 with the transcription factors CFI-1 and SOX-2 (Zhang et al. 2014; Vidal et al. 2015).

We tested whether the subtype specific IL2 markers also require the *unc-86* terminal selector, or, alternatively, whether such subtype features are regulated independently of *unc-86*-dependent ‘pan-class’ features (Zhang et al. 2014). Crossing two lateral IL2 markers, *egas-4* and *nlp-69*, and four dorsoventral IL2 markers, *egas-1, degl-2, flp-32, flp-5* into an *unc-86* null mutant allele generated by CRISPR/Cas9 genome engineering revealed that each tested marker requires *unc-86* for expression in either the lateral or dorsoventral IL2 (**Fig.1F** and **Suppl. Fig.S2B,C,D**). This observation demonstrates (1) that *unc-86/BRN3* is critical not only for controlling ‘pan-class’ features but also for IL2 subtype diversification and (2) suggests that other factors must either promote and/or restrict the ability of UNC-86 to control expression of IL-subtype-specific genes.

### *unc-39*, a SIX4/5 homolog, is continuously expressed in the lateral IL2 subtypes

Through the nervous system-wide expression analysis of all GFP-tagged homeodomain proteins, we have found that each *C. elegans* neuron class expresses a unique combination of homeodomain proteins (Reilly et al. 2020). Using a fosmid-based reporter transgene, we found that one of the homeodomain proteins, the SIX4/5 ortholog UNC-39 (Yanowitz et al. 2004), is selectively expressed in the lateral IL2, but not the dorsoventral IL2 neurons (Reilly et al. 2020). We sought to confirm this expression pattern by inserting *gfp* at the C-terminus of the *unc-39* locus by CRISPR/Cas9 genome engineering, (**Fig.2A**). Expression of the *gfp-*tagged locus, *unc-39(syb4537)*, is observed in the lateral IL2 (but not dorsoventral IL2) neurons throughout all larval and adult stages (**Fig.2A**). In the embryo, expression is also restricted to the lateral IL2 neurons and never observed in the dorsoventral IL2 neurons (Ma et al. 2021). The only other cell type that expresses the *unc-39* reporter allele throughout larval and adult stages is the AIA interneuron class. There are additional sites of expression in the embryo, consistent with *unc-39* affecting proper development of other cell types (Yanowitz et al. 2004; Lim et al. 2016). Expression of *unc-39* in the lateral IL2 neurons is eliminated in *unc-86* mutants, but not in the AIA neurons, where *unc-86* is not expressed (**Fig.2B**).

**Fig 2:**
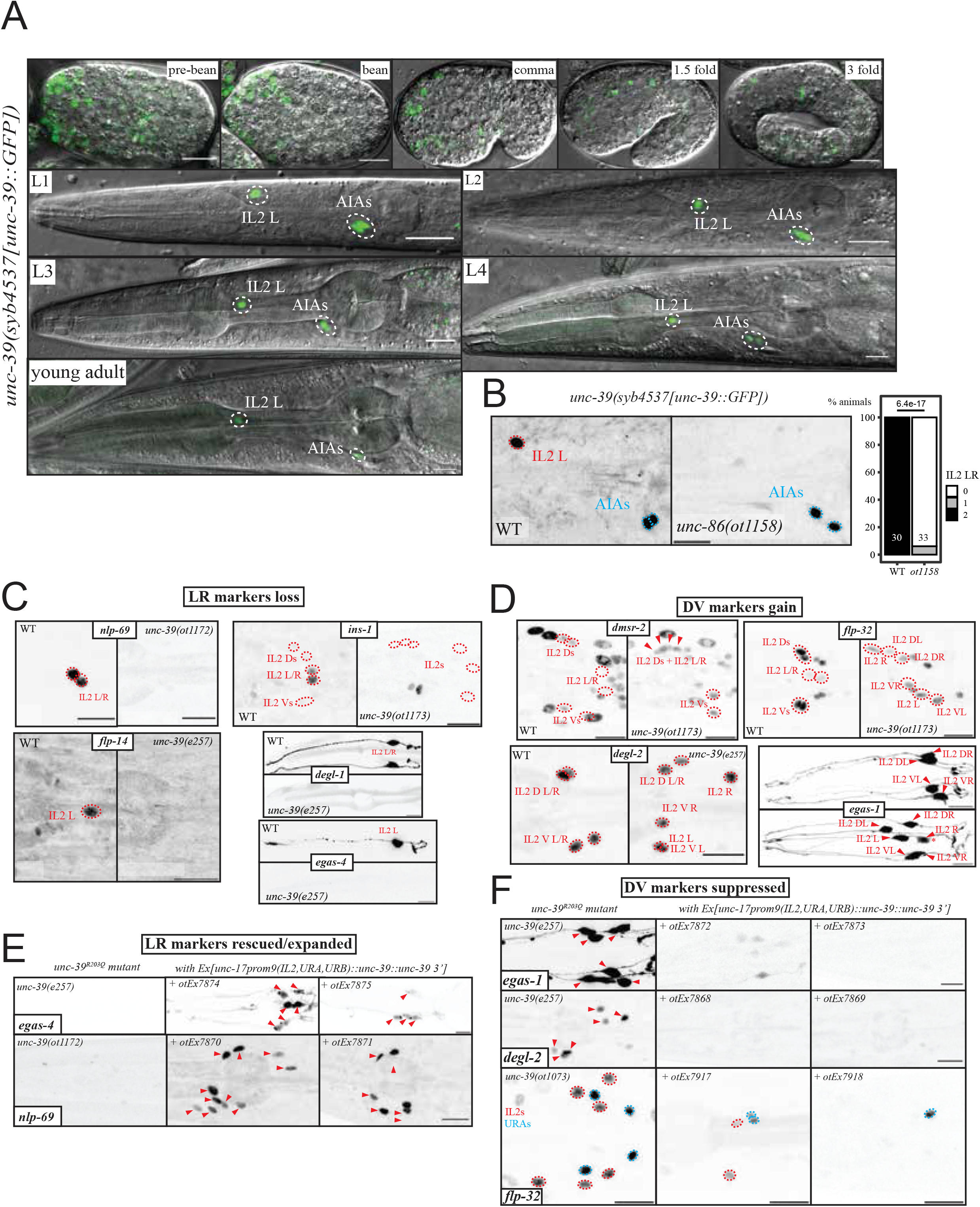
Molecular IL2 subtype diversification is controlled by *unc-39*. **A:** Expression pattern of the *unc-39(syb4537[unc-39::gfp])* reporter allele across embryonic and larval stages. **B:** *unc-39* expression in the lateral IL2 is *unc-86* dependent, with almost full loss of lateral IL2 expression in the *unc-86(ot1158)* null. AIA neurons do not express *unc-86* and *unc-39* expression is unaffected in these neurons. **C:** Lateral IL2 markers marker expression is lost in *unc-39*^R203Q^ mutant animals (either canonical *e257* allele or CRISPR/Cas9 genome engineered *ot1172* or *ot1173* alleles with identical nucleotide change). Markers are *nlp-69(syb4512[nlp-69::SL2::gfp::H2B]), ins-1(syb5452[ins-1::SL2::gfp::H2B]), flp-14(syb3323[flp-14::SL2::gfp::H2B]), degl-1 (otIs825)* and *egas-4 (otIs833)*. NeuroPAL images for cell ID and quantification is in **Suppl. Fig.4B,C**. **D**: Dorsoventral IL2 markers *dmsr-2(syb4514[dmsr-2::SL2::gfp::H2B]), flp-32(syb4374[flp-32::SL2::gfp::H2B]), degl-2(syb5229[degl-2::SL2::gfp::H2B]) and egas-1(otIs846)* are gained in all IL2 neurons of *unc-39*^R203Q^ mutant animals (either canonical *e257* allele or CRISPR/Cas9 genome engineered *ot1173* allele with identical nucleotide change). We counted gain of expression as an all or none phenotype, but the converted lateral IL2 sometimes can be dimmer (in the bottom right subpanel, *egas-1*, the asterisk marks such a case). NeuroPAL images for cell ID and quantification is in **Suppl. Fig.4B,C**. We show lateral pictures with a small rotation to show all IL2 unless precised; note that the IL2 are often displaced (**Fig.3A**) in *unc-39*^R203Q^ mutant. In the *unc-39(ok2137)* null mutant animals, the front part of the anterior ganglion moves past the anterior bulb of the pharynx (**Suppl. Fig.S4D**). **E:** Ectopic expression of *unc-39* in the IL2, URA, URB neurons in an *unc-39*^R203Q^ mutant background (*e257, ot1172* or *ot1173* allele) results in rescue of mutant phenotype in the lateral IL2 neurons and conversion of dorsal/lateral IL2 neurons. In each subpanel, the *unc-39*^R203Q^ control is shown on the left and two independent extrachromosomal array lines on the right. Two lateral IL2 markers (*nlp-69(syb4512)* rescue lines *otEx7870* and *otEx7871, egas-4/otIs833 otEx7874* and *otEx7875*,) that are lost in the hypomorph are now partially rescued and even expanded in more than just the two lateral IL2s or even the six IL2 (*nlp-69*, 10 cells in the middle panel vs 6 on the right). **F**: three dorsoventral IL2 markers (*degl-2(syb5229)* rescue lines *otEx7868 otEx7869, egas-1/otIs846* rescue lines *otEx7872* and *otEx7873* and *flp-32(syb4374)* rescue lines *otEx7917* and *otEx7918*) that are found in all the IL2 in the hypomorphic mutant animals are repressed in all subtypes upon *unc-39* misexpression. *unc-39* is able to suppress an *unc-86* dependent marker like *flp-32(syb4374)* in all IL2s but also the relatively similar URAs, which all share *unc-86* as identity regulator. In the *flp-32(ot1073*) image a ventral view is provided. Quantification is in **Suppl. Fig.S4E,F**.

Continuous expression of a transcription factor throughout the life of a cell is often an indication of autoregulation. We tested whether *unc-39* autoregulates its own expression by first defining *cis-*regulatory elements required for continuous expression in the IL2 and AIA neurons. We found that neuronal *unc-39* expression in the lateral IL2 neurons and the AIA neurons is recapitulated by a 2.3 kb region directly upstream of the *unc-39* locus (**Suppl. Fig.S3A**). Expression of this promoter fusion construct is downregulated in the *unc-39* mutant animals (**Suppl. Fig.S3A**). Mutation of a small region with a cluster of several predicted homeodomain binding sites within this reporter constructs affected its expression in both lateral IL2 and AIA neurons (**Suppl. Fig.S3B**). We introduced this mutation in the endogenous *unc-39(syb4537)* reporter allele using CRISPR/Cas9 genome engineering, and also observed loss of expression in both IL2 and to a lesser extent in AIA (**Suppl. Fig.S3C**). This regulatory mutation is not the result of a failure to initiate *unc-39* expression in IL2 and AIA since these mutant animals display wild-type-like expression of the *gfp*-tagged *unc-39* locus until the first larval stage, after which expression fades away (**Suppl. Fig.S3C**). Hence, *unc-39* expression is maintained through a mechanism that is independent of its embryonic initiation. Since maintained expression genetically depends on *unc-39*, as well as on predicted homeodomain binding sites in its promoter, we surmise that *unc-39* autoregulates its expression.

### *unc-39* is required for IL2 subtype diversification

To investigate the functional effects of *unc-39*, we relied mainly on the canonical *unc-39(e257)* loss of function allele, which contains a R203Q mutation in the SIX domain of *unc-39* (Yanowitz et al. 2004). To avoid linkage issues with many markers, we introduced the same mutation in some of our reporter strains. We find that *unc-39*^R203Q^ mutant animals display a loss of the expression of all five tested lateral specific markers, *egas-4, degl-1, ins-1, flp-14* and *nlp-69* (**Fig.2C**). Conversely, *unc-39*^R203Q^ mutant animals display a concomitant gain of all four tested dorsoventral IL2 marker expression (*egas-1, flp-32, dmsr-2, degl-2*) in the lateral IL2 neurons of *unc-39* mutants (**Fig.2D**), indicating that the lateral IL2 neurons have adopted the identity of the dorsoventral IL2 neurons.

*unc-39* selectively affects IL2 subtype specification, since we observe no effects on the expression of identity features that are expressed by all six IL2 neurons. Specifically, expression of the vesicular transporter *unc-17* and the kinesin *klp-6*, both controlled by *unc-86* in all six IL2 neurons (Zhang et al. 2014), is not affected in *unc-39(e257)* mutant animals or the *unc-39(ok2137)* null (**Suppl. Fig.S4D**).

### *unc-39* is sufficient to transform IL2 subtype identities

To ask whether *unc-39* is not only required but also sufficient to induce lateral IL2 subtype identity and to suppress dorsoventral IL2 identity, we ectopically expressed *unc-39* in the dorsoventral IL2 neurons, using a pan-IL2 driver, an enhancer fragment from the *unc-17* locus (Serrano-Saiz et al. 2020). We verified that this driver is unaffected in *unc-39* null and hypomorphic animals (**Suppl. Fig.S4D**). In an *unc-39(e257)* background, this transgene was not only able to rescue the loss of the lateral IL2 marker genes *egas-4* and *nlp-69* in the lateral IL2 neuron, but the transgene was also able to induce *egas-4* and *nlp-69* expression in the dorsoventral IL2 subtype (**Fig.2E**). Conversely, ectopic expression of the dorsoventral IL2 markers in the lateral IL2 neurons, observed in *unc-39* mutants, is not only suppressed by this transgene, but expression of these dorsoventral IL2 subtype markers are repressed in the dorsoventral IL2 subtypes as well (**Fig.2F**). Together, these findings demonstrate that *unc-39* is not only required, but also sufficient to promote lateral IL2 identities.

In animals that express *unc-39* under the *unc-17prom9* driver, we also noted occasional ectopic expression of the lateral IL2 marker *nlp-69* in a few additional neurons, which are likely the URA and URB neurons, which also express the *unc-17prom9* driver (Serrano-Saiz et al. 2020). *unc-86* is known to work as a putative terminal selector in the URA and URB neurons (Zhang et al. 2014). We therefore surmise that *unc-39*, in conjunction with *unc-86* is able to induce IL2 lateral identity in other cellular contexts as well. Vice versa, ectopic *unc-39* expression is also sufficient to suppress dorsoventral IL2 marker genes that normally happen to be expressed in other neurons. Specifically, the neuropeptide-encoding *flp-32* marker is normally strongly expressed in the *unc-86-*dependent URA and dorsoventral IL2, and dimly in the lateral IL2 neurons. Ectopic *unc-39* expression with the *unc-17prom9* driver is able to almost fully suppress *flp-32* expression in these neurons (**Fig.2F)**, to an even lower levels than the already weak wild type lateral IL2 expression **(Fig.1E)**. Taken together *unc-39* is sufficient to activate and to repress specific target genes not just in the context of the IL2 neurons, but also in other cellular contexts that normally require the *unc-86* homeobox gene for their proper differentiation.

### Transformation of other phenotypic features of IL2 subtypes in *unc-39* mutants

A transformation of lateral IL2 to dorsoventral IL2 identity in *unc-39* mutants is not only indicated by molecular markers, but is also further supported by an analysis of IL2 morphology. First, we note that in *unc-39(e257)* mutants, lateral IL2 cell bodies can become closely associated with dorsoventral IL2 cell bodies and their dendritic projections are often fasciculated with the dorsoventral IL2 dendrites (**Fig.3A, Suppl. Fig.S5A**). Moreover, the positional stereotypy of the lateral IL2 soma is transformed to the much more variable dorsoventral IL2 soma position that we had previous described (Yemini et al. 2021).

We serendipitously discovered another subcellular feature that distinguishes lateral from dorsoventral IL2 neurons: all the IL2 localize the synaptic active zone marker CLA-1/Clarinet at the dendritic endings in the nose of the worm, but the lateral IL2 neurons also show CLA-1 punctae localized along the dendrite (**Fig.3B**). Synaptic vesicle machinery had previously been observed in ciliary endings of other sensory neurons (Mazelova et al. 2009; Datta et al. 2015; Ojeda Naharros et al. 2017), but had not been examined in the IL2 neurons to date. Dendritic CLA-1 localization is likely unrelated to EV release by IL2 neurons (Wang et al. 2014), since we find it to be unaffected in *klp-6(ky611)* mutants, which lack the kinesin that transports EVs. In *unc-39(e257)* mutants, CLA-1 localization along the length of the dendrite is reduced if not eliminated, comparable to what is normally seen only in the dorsoventral IL2 subtypes in wild-type animals. Conversely, ectopic *unc-39* expression in all six IL2 neurons results in CLA-1 signals being observed along the length of all IL2 dendrites, comparable to what is normally only observed in the lateral IL2 neurons (**Fig.3B**). We conclude that CLA-1 dendritic localization is differentially regulated in the two IL2 subtypes and that *unc-39* controls this feature as well.

Upon entry into the dauer stage, the six IL2 neurons adopt further subtype-specific features. On a morphological level, the dorsoventral, but not lateral IL2 neurons, develop extensive dendritic branches (Schroeder et al. 2013)(**Fig.3C**). *unc-39(e257)* animals form SDS-resistant dauer normally, but the lateral IL2 neurons now seem to display dendritic branches, i.e. adopt features normally exclusive to the dorsoventral IL2 neurons (**Fig.3C**). These branches appear to merge with the IL2 dorsoventral branches. Conversely, ectopic expression of *unc-39* in all six IL2 neurons almost completely suppresses branch formation in the dorsoventral IL2 neuron, making them resemble the branchless lateral IL2 neurons.

**Fig 3:**
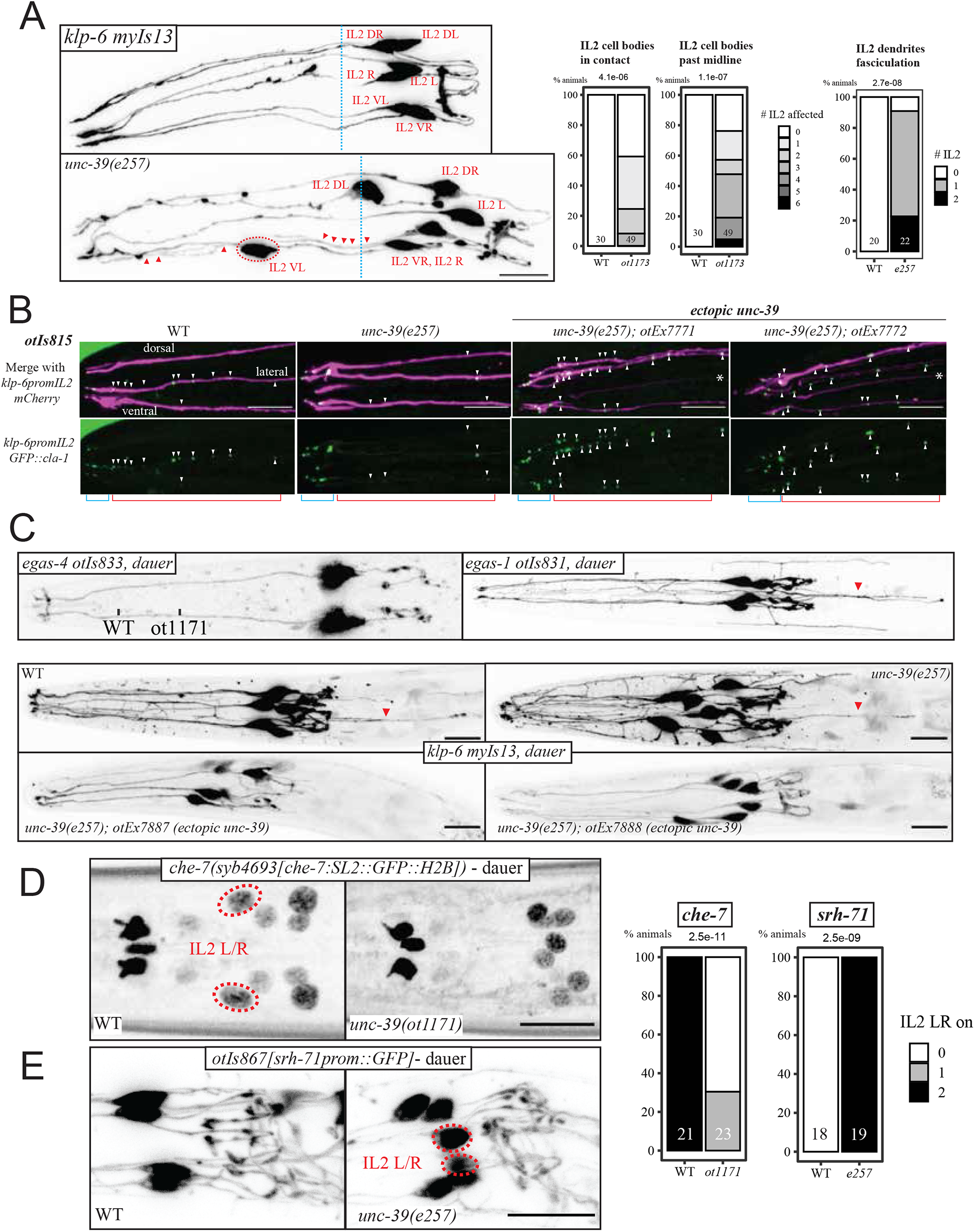
Morphological IL2 subtype diversification is controlled by *unc-39*. **A:** The neurites of all six IL2 are labelled by the *unc-39* independent marker *myIs13[klp-6prom::GFP]*. Left panel: In wild-type animals (WT), IL2 cell bodies are usually adjacent to the anterior bulb of the pharynx and their processes run in six fascicles alongside, as shown with the *klp-6 (myIs13)* cytoplasmic reporter. In the *unc-39* hypomorph, the lateral IL2 neurons are often out of position, fully overtaking the midline of the anterior bulb (red circled neuron past the blue dash line). Their dendrites join the the dorsal or ventral tracts (red triangles, see depth-map **Suppl. Fig.S4A**). Right: we quantified neurite fasciculation defects (abnormally close tracts). We also considered cell body position defect like anteriorly displaced IL2 (crossing the blue midline) or IL2 nuclei entering into contact with each other, using NeuroPAL *otIs669* wild type and *unc-39*^R203Q^.. **B:** A *cla-1* reporter transgene, that uses *klp-6* to label the neurite with mCherry and expresses a GFP-tagged short *cla-1* isoform (*otIs815*) is found in the endings of all six IL2 dendrites (blue segment), but also further along the dendrites (red segment; punctae marked with white triangles) of the lateral IL2 neurons. This second group punctae (along the dendrite) are eliminated in the *unc-39*^R203Q^ hypomorph, while upon misexpression of *unc-39* they frequently appear in multiple dendrites. See quantifications in **Suppl. Fig.S5B**. **C:** Dauer arborization changes as shown by *myIs13[klp-6prom::GFP];* worms are SDS selected. **Top:** In wild-type animals, IL2 neurons display dauer specific arborization, with increased branching in the dorsoventral subtypes compared to the lateral subtypes. We show dauer pictures of our subtype specific intregrated cytoplasmic reporter, *egas-4 (otIs833)* top left in ventral position, *egas-1/otIs831* top right. **Bottom:** IL2 branching imaged with *klp-6 (myIs13)* cytoplasmic reporter shows increased branching of the lateral IL2 neurons of *unc-39(e257)*, which is difficult to quantify, while overexpressing *unc-39* using the *unc-17prom::gfp* driver in all IL2 neurons (transgenic lines *otEx7887* and *otEx7888)* strongly suppress this arborization.The backward projecting neurites we saw are labelled with red triangles. See quantifications in **Suppl. Fig.S5C**. **D:** The innexin *che-7* reporter allele *syb4693* is gained in dauer in the lateral IL2 in WT worms, but not in *unc-39*^R203Q^ mutants. **E:** The *srh-71* GPCR reporter transgene (*otIs867*) is expressed in all IL2 in wild-type worms, and lost from the lateral IL2 when they go into dauer. In *unc-39*^R203Q^ mutant dauer, expression remains in the lateral IL2 neurons while expression is eliminated in wild-type animals.

We have also noticed that in the dauer stage animals dorsoventral IL2 neurons appears to project neurites posterior to the cell body, as visualized with the cytoplasmic and subtype specific *egas-1* and *egas-4* markers (**Fig.3C**). While those projections are hard to score in *unc-39* mutant animals, they obviously disappear upon ectopic expression of *unc-39* in all IL2 neurons. This observation further corroborates the notion that *unc-39* controls subtype-specific morphological features of the IL2 neurons (**Fig.3C**).

Entry into the dauer stage also leads to IL2 subtype-specific changes in gene expression. These include the dauer-specific induction of a GFP-tagged electrical synapse protein, CHE-7, exclusively in the lateral IL2 neurons (Bhattacharya et al. 2019). Conversely, the GPCR encoding gene *srh-71*, which, under replete conditions, is expressed in all six IL2 neurons, becomes selectively repressed in the lateral IL2 neurons (Vidal et al. 2018). Both of these molecular remodeling events fail to occur in *unc-39* mutant dauers (**Fig.3D,E**). A *che-7* transcriptional CRISPR reporter fails to be properly induced in lateral IL2 neurons and, conversely, a *srh-71* reporter construct is derepressed in the lateral IL2 neurons of *unc-39* mutant dauer animals (**Fig.3E**). After dauer recovery, *srh-71* is turned off in wild-type animals, while in *unc-39* mutant animals, *srh-71* expression persists (**Suppl. Fig.S5D)**. Taken together, both molecular as well as morphological data is consistent with what can be referred to as a ‘homeotic identity transformation’ of the lateral IL2 to the dorsoventral IL2 neurons in *unc-39* mutants.

### Nested homeobox gene function also controls subtype diversification in the RMD neck motor neuron class

Our functional analysis of the IL2 neurons reveal a nested set of homeobox gene function: A homeobox terminal selector controls all identity features of a neuron and a subordinate, subtype-specific homeobox gene differentiates the identity of neuronal subtypes. Does the concept of nested homeobox gene function apply in other neuron classes as well? Intriguingly, our genome- and nervous-system wide expression pattern analysis of homeodomain proteins revealed that another member of SIX homeodomain protein family, the SIX3/6-like CEH-32 protein, is expressed in a subtype-specific manner in yet another radially symmetric class of neurons, the RMD neck motorneurons (Reilly et al. 2020)(**Fig.4A**). A CRISPR/Cas9-engineered *gfp* reporter allele is expressed throughout all larval and adult stages only in the dorsoventral RMD neuron subclass, not the lateral RMD subclass **(Fig.4B)**.

**Fig 4:**
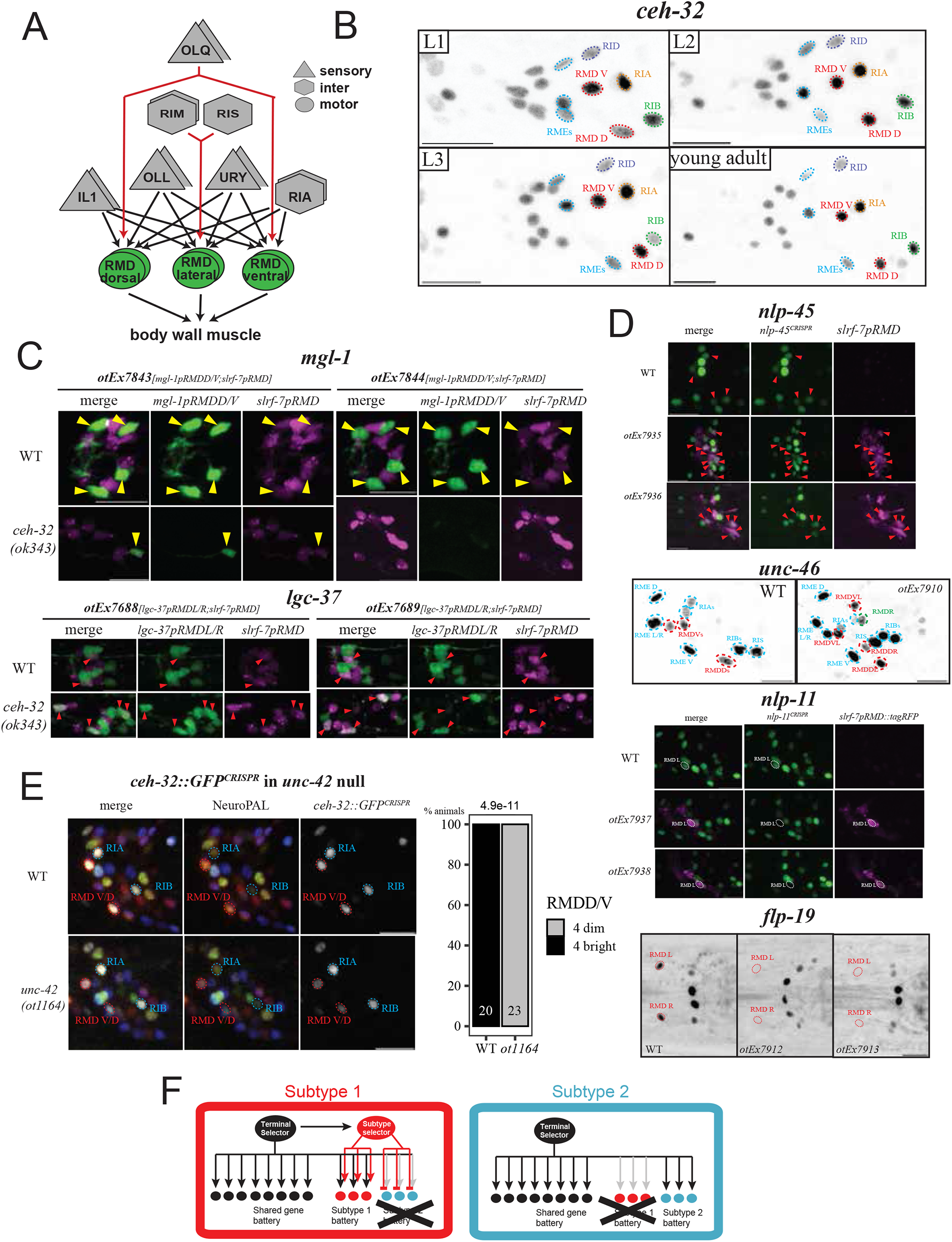
Another SIX homeobox gene diversifies another radially symmetric neuron class. **A:** RMD neck motor neuron overview. Other connectivity difference (e.g. dopamine neurons; or lateral neuron specific feedback to RIA) not shown. **B:** A GFP tagged *ceh-32* reporter allele *(ot1040)* is expressed across all larval stages in the ventral and dorsal RMD subtypes (circled in red), located respectively dorsally and ventrally. *ceh-32* expression is never observed in the lateral RMD. The *ceh-32* expressing neuron classes that are closest to the RMD (RIA/RIB/RID/RME) are labelled. **C:** Loss of function experiments. RMD subtype marker expression in *ceh-32(ok343)* null mutant animals. *slrf-7*^*prom*^*::TagRFP* is present on all transgenes to label overall RMD fate. Top, expression of a *mgl-1* reporter, expressed in dorsoventral RMD (yellow triangles), is lost in two different lines (*otEx7843* and *otEx7844)*. Bottom, a *lgc-37* reporter, expressed in lateral RMD (red triangles) is ectopically expressed in more than just two RMDs, as seen in two different lines (*otEx7688* and *otEx7689)*. The number of *slrf-7(+)* neurons is not affected (expression in the dorsoventral RMDs appears brighter in *ceh-32* mutants). Quantifications are found on **Suppl. Fig.S6C** **D:** Gain of function experiments. *ceh-32* is misexpressed in all six RMD neurons using the *slrf-7* driver. The dorsoventral subtype markers *nlp-45* (reporter allele *ot1032)* and *unc-46* (fosmid *otIs568)* can be found in more than the usual four expected RMDs (using *ceh-32* misexpressing arrays *otEx7935* and *otEx7936* for *nlp-45* reporter allele, and *otEx7909* for *unc-46* fosmid line; another line, *otEx7910*, had no effect). In the case of *nlp-45*, the cells expressing both *nlp-45* and a *slrf-7p::tagRFP* marker as part of the *otEx7935/7936* misexpression arrays are shown with red triangles. The lateral RMD subtype markers *nlp-11* (reporter allele *syb4759)* and *flp-19 (*reporter allele *syb3278)* are repressed by pan-RMD expression of *ceh-32 (*expressed with arrays *otEx7937* and *otEx7938* in the *nlp-11* reporter allele and *otEx7912* and *otEx7913* in the *flp-19* reporter allele). In the *nlp-11* reporter allele background, we also used *slrf-7p::tagRFP* as part of the misexpression array to help identifying the RMD neurons. The lateral RMDs are circled in white. For *flp-19*, we show ventral images, the lateral RMD are circled in red and would be anterior to the two bright cells in the middle (AIAs). Quantifications are found on **Suppl. Fig.S6D**. **E:** In an *unc-42(ot1164)* null allele, the *ceh-32* reporter allele *ot1040* becomes dimmer in both dorsal and ventral RMDs. We quantified the RMDD/V (circled red) expression as bright vs dim, using the brightness of RIA and RIB (circled blue) as reference. Those neurons do not express *unc-42* and stay unchanged in *unc-42* null mutant animals. p-values are done via Fisher two-sided test. **F:** Summary. Terminal selectors operate in a nested regulatory configuration in which they regulate the expression of subtype terminal selectors that specify subtype identity. Loss of a subtype selector results in homeotic identity transformation to another subtype identity. Subtype selectors either promote the ability of a terminal selector to activate specific subtype identity features (‘subtype 1 battery’) or antagonize the ability of a terminal selector to activate specific subtype identity features (‘subtype 2 battery’). Open questions are how the subtype-specificity of subtype regulators are controlled and whether subtype selectors act directly on target genes (as indicated here), or through intermediary factors (e.g. by activating a repressor to suppress alternative subtype fates).

The RMD neck motor neuron class is entirely distinct from the IL2 sensory neuron class, not only in terms of overall location in a different ganglion, function (motor vs. sensory neuron), but also morphology (i.e. neurite projection patterns), synaptic connectivity and molecular composition. But like the IL2 neuron class, the RMD class is also composed of three bilateral neuron pairs, a dorsal, lateral and ventral pair that share a plethora of anatomical, as well as molecular features among each other (White et al. 1986; Hobert et al. 2016; Taylor et al. 2021)(**Fig.4A**). Similar to the IL2 neuron case, all six RMD neurons are genetically specified by a homeobox-type terminal selector gene, the Prop1-like *unc-42* gene, which controls the expression of essentially all known terminal identity features of the RMD neurons (Berghoff et al. 2021). Yet, in spite of many similarities, the dorsoventral and the lateral pairs do show differences in synaptic connectivity. as well as molecular composition (White et al. 1986; Hobert et al. 2016; Taylor et al. 2021)(**Fig.4A**). For example, the metabotropic glutamate receptor *mgl-1* has been found to be expressed in the dorsoventral RMD pairs, but not the lateral pair (Greer et al. 2008; Zhang et al. 2014), while the GABA receptor subunit *lgc-37* is selective expressed in the lateral, but not dorsoventral RMD pairs (Greer et al. 2008; Zhang et al. 2014; Gendrel et al. 2016; Taylor et al. 2021). Through the CRISPR/Cas9-mediated engineering of reporter alleles, we validated other genes that scRNA analysis found to be subtype-specifically expressed, including several neuropeptide-encoding genes (*flp-19, nlp-11, nlp-45*)(**Fig.4D**).

Armed with these markers of subtype identity, we asked whether CEH-32 parallels the subtype diversification function of UNC-39 in the context of the RMD neurons. We found that expression of a reporter transgenes that monitors expression of *mgl-1* in the dorsoventral RMD neuron is lost in *ceh-32* mutant animals (**Fig.4C, Suppl. Fig.S6A**). Conversely, the *lgc-37* gene, normally restricted to the lateral RMD neurons, becomes derepressed in the dorsoventral RMD neurons (**Fig.4C, Suppl. Fig.S6A**). In contrast, *ceh-32* does not affect expression of a pan-RMD marker, *C42D4*.*1/srlf-7* (**Fig.4C; Suppl. Fig.S7**).

To test for sufficiency of *ceh-32* function, we expressed *ceh-32* in all six RMD neurons using the promoter of the pan-RMD *C42D4*.*1/slrf-7* promoter. Such transgenic animals showed gain of several dorsoventral subtype markers in the lateral RMD neurons and a concomitant loss of left/right subtype markers (**Fig.4D, Suppl. S6B**). Taken together, our data indicates a homeotic identity change of RMD subtypes that is akin to what we observed in the IL2 neurons upon manipulation of *unc-39* expression.

The pan-RMD terminal selector *unc-42* affects both the subtype-specific markers, as well as the pan-RMD markers (Gendrel et al. 2016; Berghoff et al. 2021; Sun and Hobert 2021)(**Suppl. Fig.S8**). Moreover, *ceh-32* expression in the dorsoventral RMD neurons in diminished in *unc-42* mutants (**Fig.4E**). Hence, as in the case of the IL2 neurons, a subtype-specific SIX homeodomain protein controls their subtype-specific identity features, while a pan-class homeobox terminal selector controls both pan-class identity features as well as subtype-specific features.

## DISCUSSION

An analysis of the taxonomic relationship between neuronal cell types in many different animal brains reveals closely related neuronal subtypes at the terminal branch point of such taxonomies that share a multitude of anatomical and molecular features but differ in a select number of phenotypic properties. In theory, one can envision that such very closely related cell types are independently specified by distinct regulatory programs. In the cases that we describe here, this is not the case. We rather uncover, in two distinct cellular contexts (IL2 neuron class and RMD neuron class), a nested regulatory architecture that specifies differences between closely related neuronal cell types (**Fig.4F**). Shared, as well as subtype-specific features of a neuronal cell type are controlled by a terminal selector transcription factor that is expressed in all subtypes (e.g. UNC-86 in all IL2 subtypes and UNC-42 in all RMD subtypes)(**Fig.4F**). Subtype-specificity of select identity features are dictated by ‘subtype terminal selectors’ that modulate the ability of terminal selector to control the expression of specific target genes (**Fig.4F**). Depending on the target genes, a subtype terminal selector like UNC-39 may either assist the ability of a class-specific terminal selector like UNC-86, to bind DNA and/or to activate transcription of subtype-specific features (e.g. lateral IL2 markers), or, on other target genes, prevent the terminal selector from doing so (e.g. dorsoventral IL2 markers)(**Fig.4F**). Given that the diversification of neuronal cell types into closely related subtype is observed in many different contexts not only in invertebrate but also vertebrate brains (Shekhar et al. 2016; Cembrowski and Spruston 2019; Network 2021; Yao et al. 2021), we envision the nested gene regulatory principle described here to be broadly applicable, with similarities of two cell types being defined by the same terminal selector, but diversified by subordinate transcription factors (‘subtype terminal selectors’).

Our study confirms the importance of homeobox genes in neuronal identity control (Reilly et al. 2020). The mutant phenotype that we describe here are conceptually reminiscent of body segment identity transformation in HOX cluster mutants. In fact, we had previously described that *C. elegans* HOX genes also act in terminal motor neuron differentiation to diversify motor neuron subtype identities along the anterior/posterior axis (Kratsios et al. 2017). Viewing neuronal subtype identity transformations as ‘homeotic’ places them into the rich framework of evolutionary thought that started with William Bateson ‘s definition of homeosis in 1894 (Bateson 1894; Arlotta and Hobert 2015). While it remains debatable whether homeotic identity changes of entire body parts have been drivers of evolution (Akam 1998), homeotic identity transformations of smaller units, i.e. individual cell types, can be more easily envisioned to have played a role in evolution of the composition and complexity of tissue types (Arlotta and Hobert 2015). Specifically, it is conceivable that neuronal subtypes evolved from a homogeneous, terminal selector-controlled state in which a group of neurons all shared the same features. The gain of expression of a subtype selector in a subgroup of these cells may have then enabled to modulate the expression of a subset of the shared identity features of this group of neurons.

## MATERIAL AND METHODS

### Strains and mutant alleles

A complete list of mutant and transgenic strains can be found in **Suppl. Table S1**. Due to linkage issues (*nlp-69, che-7*, NeuroPAL integrant *otIs669*), we regenerated the *e257* allele, a G>A mutation that alters arginine 203 to glutamine in the SIX domain in several different strain backgrounds, using CRISPR/Cas9 genome engineering following (Dokshin et al. 2018). We use the following repair template (bold: sgRNA, italics: G>A mutation): CGTGGCAAAGAAC**TGAATCCAGTGGAAAAATAT**C*a*GCTGAGACGAAAGTTTCCGGCTC CGAAAACAATTTGG. The G>A edit eliminates the PAM site, the edited allele is identical to *e257*.

The *unc-39* binding site mutant *ot1193* was generated via the same CRISPR protocol, with the guide CC939 and repair template shown in **Suppl. Fig.S3B**.

We generated in the background of the *ceh-32* reporter allele *ot1040* a novel *unc-42* null allele, *unc-42(ot1164)*, using the same CRISPR protocol as described above, deleting the entire coding gene of *unc-42*. We used the two following guides, GCTCATtgtgtgagtgaaag and tctcactgatagactaatgt, and the following repair oligo: ttcggtcacccctcactttccacatttctccgctttgttggtacgtacatttacaaaaagacaacggttt.

Mixes use the described amount, were injected as simple arrays except for the *ceh-32* and *unc-39* rescue strains which were injected as complex. pBluescript DNA as an injection filler to reach 100ng/ul for simple injection mixes, and OP50 genomic DNA for complex mixes. Integration was done via gamma radiation, with the strains being outcrossed at least 3x.

The coinjection marker *inx-19promAVG::TagRFP* is from (Oren-Suissa et al. 2016). The *inx-6prom(2TAAT-deletion)::GFP* coinjection marker is from (Bhattacharya and Hobert 2019) and labels the procorpus of the pharynx.

### Reporter construction

Reporter alleles for *che-7, dmsr-2, degl-2, flp-5,flp-14, flp-19, flp-32, ins-1, nlp-69, nlp-11*, (see strain list) were generated using CRISPR/Cas9, inserting a SL2::gfp::h2b cassette at the C-terminus of the respective gene. These strains, as well as the *unc-39* reporter allele, *syb4537*, a direct fusion of *gfp* to the C-terminus of *unc-39*, were generated by Sunybiotech. For *srh-71*, we reinjected the *srh-71* PCR fusion mix from (Vidal et al. 2015), yielding an *otEx7765 pha-1* rescue line that was integrated.

Promoter fusions for the *egas1, egas-4, degl-1* and *unc-39* genes were Gibson-cloned into the pPD95.75 backbone, using either the SphI/XmaI multiple cloning sites for promoters and KpnI/ApaI for coding sequence/ 3 ‘ UTR. Primer sequences are shown below:

- ***egas-1*** 1422bp promoter: **fwd** aaccagtgtaccgtccatctg **rev** atgatttataaggacgttagggaagtttg
- ***egas-4*** 278bp promoter: **fwd** caaatcaaaaacaggtctcatgtacatac **rev** ttcttgactgaatcctaataggaaattc
- ***degl-1*** 230bp promoter: **fwd** aacaacgctaacaagtaaccagatg **rev** aaaatttacaaaacactgagcttgtgc
- ***slrf-7*** 570bp promoter: **fwd** gattcgcggaactctagtaaattgaatc **rev** cctgaaatcaaccatttccatcaagaag
- ***unc-39*** 2260bp promoter: **fwd** gaagagtgtcgaatattgcagcag **rev** tggaatactactcatcttgcaaacgttc
- ***unc-39*** 1754bp **CDS and 3 ‘ UTR**: **fwd** ATGACAGACCATCCGCCAATTG **rev** tgccaacattgattgatctctacg
- ***ceh-32*** 3694bp **CDS and 3 ‘ UTR: fwd** atgttcactccagaacagttcac **rev** tattttgcagattcgcattacttcgg

### Misexpression approaches

The *unc-39* and *ceh-32* CDS and 3 ‘ UTR fragment described above were misexpressed in all six IL2 or RMD neurons, respectively, using the *unc17prom9* driver for the IL2 neurons (Serrano-Saiz et al. 2020) and, for the RMD neurons, a 570 bp of 5 ‘ region of the C42D4.1/srlf-7 gene, which encodes a small secreted protein that is a member of a large family of SXP/RAL2 related proteins, characterized by an ANIS5 Pfam domain (PF02520). scRNA analysis had shown expression in the RMD neurons (Taylor et al. 2021) and this pattern was confirmed with promoter fusions (Lorenzo et al. 2020). We find that 570bp of the C42D4.1/ srlf-7 promoter drive expression of TagRFP or GFP in all six RMD neurons, with occasional expression in the SAA neurons (**Suppl. Fig.S2**).

### Microscopy and image processing

Worms were anesthetized using 100 mM sodium azide (NaN3) and mounted on 5% agarose pads on glass slides. Z-stack images (40x for L4 animals .7 micron steps, 63x for L1 with .4 steps) were acquired using a Zeiss confocal microscope (LSM880) or Zeiss compound microscope (Imager Z2) using the ZEN software. Maximum intensity projections of 2 to 30 slices were generated with the ImageJ software (Schindelin et al. 2015). All scale bars shown in this paper are 10 microns. We show our reporters as inverted gray, with the same range between genotypes.

In the case of NeuroPAL images, we split the channels as the GFP signal channel in gray, and the NeuroPAL coloring channels (mTabBFP2,CyOFP,mNeptune). We adjusted when needed the gamma to 0.5 but for the latter group of channels only. Images are shown as GFP signal in inverted gray in main figures, and for each of those the merge / GFP in gray / Neuropal colors are shown in supplements. Depth coloring for **Suppl. Fig.S4A** was done in Fiji via the Image/Hyperstacks/Temporal color-code command.

In the case of *dauer* experiments, we grew gravid adults for 6-8 days at 25C, washed the plates with 1% SDS into multiple tubes and left the worms to incubate under gentle agitation for 20 minutes at least and 2 hours at most. Before imaging, we washed off worms in M9 one plate at a time and replated the living dauer and non-dauer carcasses on an uncoated plate, imaging for one hour or so before switching to a new tube of worms in SDS.

### Data analysis and statistics

All statistics and plots were done in R. We used the R tidyverse package collection and the ggplot2 graph library. We used conda to manage packages, and Jupyter notebooks via the ‘r-irkernel’ package. The notebooks and a conda manifest file are available on request. We created bar plots with ‘geom_bar’, organized scoring data into contingency tables and used two-sided Fisher test for p-values for most experiments except for misexpression experiments. In those case we plotted neuron counts with ‘geom_beeswarm’, and used a Wilcoxon rank sum test to compare each misexpression line to the relevant control (*unc-39* hypomorph, *ceh-32* WT). We adjusted p-values for multiple testing via Holm ‘s method. We did not determine sample sizes in advance, but used as baseline 30-40 worms when scoring on a dissecting scope, and for confocal / microscope scoring we stopped at 10-15 worms for fully penetrant phenotypes and 15-25 otherwise. Extrachromosomal lines were scored as 10-15 worms, with a control and at least 2 lines. On the graphs, the number on each bar each show the number of worms scored per genotype.

## ACKNOWLEDGEMENTS

We thank Chi Chen for generating transgenic lines, Tessa Tekieli for an *unc-86* null allele and pictures of *flp-5* crossed to it, Maryam Majeed for generating the IL2 *cla-1* line integrated here and a *flp-*3 IL1 reporter, Robert Fernandez for providing multiples neuropeptide reporter alleles already crossed to NeuroPAL, Molly Reilly for generating the *ceh-32* fosmid rescue, Eduardo Leyva-Dííaz for *unc-42* CRISPR deletion allele reagents and comments on the manuscript, Surojit Sural for help and advice on dauer experiments and Berta Vidal for a *srh-71* PCR fusion injection mix. We thank the Hobert lab members and Nathan Schroeder for advice and feedback. Some strains were provided by the CGC, which is funded by NIH Office of Research Infrastructure Programs (P40 OD010440). This work was funded by NIH R21NS106843 and the Howard Hughes Medical Institute.

## REFERENCES

Akam M. 1998. Hox genes, homeosis and the evolution of segment identity: no need for hopeless monsters. Int J Dev Biol 42: 445–451.

Al-Sheikh U, Kang L. 2020. Mechano-gated channels in C. elegans. J Neurogenet 34: 363–368.

Arendt D, Bertucci PY, Achim K, Musser JM. 2019. Evolution of neuronal types and families. Curr Opin Neurobiol 56: 144–152.

Arlotta P, Hobert O. 2015. Homeotic Transformations of Neuronal Cell Identities. Trends in neurosciences 38: 751–762.

Bateson W. 1894. Materials for the study of variation, treated with especial regard to discontinuity in the origin of species. Macmillan, London.

Berghoff EG, Glenwinkel L, Bhattacharya A, Sun H, Varol E, Mohammadi N, Antone A, Feng Y, Nguyen K, Cook SJ et al. 2021. The Prop1-like homeobox gene unc-42 specifies the identity of synaptically connected neurons. eLife 10.

Bhattacharya A, Aghayeva U, Berghoff EG, Hobert O. 2019. Plasticity of the Electrical Connectome of C. elegans. Cell 176: 1174–1189 e1116.

Bhattacharya A, Hobert O. 2019. A new anterior pharyngeal region specific fluorescent co-transformation marker. MicroPubl Biol 2019.

Cajal Ry. 1911. Histologie du système nerveux de l’homme et des vertébrés. Maloine, Paris.

Cembrowski MS, Spruston N. 2019. Heterogeneity within classical cell types is the rule: lessons from hippocampal pyramidal neurons. Nat Rev Neurosci 20: 193–204.

Cook SJ, Jarrell TA, Brittin CA, Wang Y, Bloniarz AE, Yakovlev MA, Nguyen KCQ, Tang LT, Bayer EA, Duerr JS et al. 2019. Whole-animal connectomes of both Caenorhabditis elegans sexes. Nature 571: 63–71.

Datta P, Allamargot C, Hudson JS, Andersen EK, Bhattarai S, Drack AV, Sheffield VC, Seo S. 2015. Accumulation of non-outer segment proteins in the outer segment underlies photoreceptor degeneration in Bardet-Biedl syndrome. Proc Natl Acad Sci U S A 112: E4400–4409.

Dokshin GA, Ghanta KS, Piscopo KM, Mello CC. 2018. Robust Genome Editing with Short Single-Stranded and Long, Partially Single-Stranded DNA Donors in Caenorhabditis elegans. Genetics 210: 781–787.

Finney M, Ruvkun G. 1990. The unc-86 gene product couples cell lineage and cell identity in C. elegans. Cell 63: 895–905.

Gendrel M, Atlas EG, Hobert O. 2016. A cellular and regulatory map of the GABAergic nervous system of C. elegans. eLife 5.

Greer ER, Perez CL, Van Gilst MR, Lee BH, Ashrafi K. 2008. Neural and molecular dissection of a C. elegans sensory circuit that regulates fat and feeding. Cell Metab 8: 118–131.

Hall DH, Altun Z. 2007. C. Elegans Atlas. Cold Spring Harbor Laboratory Press.

Hobert O. 2013. The neuronal genome of Caenorhabditis elegans. WormBook: 1–106.

Hobert O, Glenwinkel L, White J. 2016. Revisiting Neuronal Cell Type Classification in Caenorhabditis elegans. Curr Biol 26: R1197–R1203.

Kratsios P, Kerk SY, Catela C, Liang J, Vidal B, Bayer EA, Feng W, De La Cruz ED, Croci L, Consalez GG et al. 2017. An intersectional gene regulatory strategy defines subclass diversity of C. elegans motor neurons. eLife 6.

Lee H, Choi MK, Lee D, Kim HS, Hwang H, Kim H, Park S, Paik YK, Lee J. 2012. Nictation, a dispersal behavior of the nematode Caenorhabditis elegans, is regulated by IL2 neurons. Nat Neurosci 15: 107–112.

Leyva-Diaz E, Masoudi N, Serrano-Saiz E, Glenwinkel L, Hobert O. 2020. Brn3/POU-IV-type POU homeobox genes-Paradigmatic regulators of neuronal identity across phylogeny. Wiley interdisciplinary reviews Developmental biology: e374.

Lim MA, Chitturi J, Laskova V, Meng J, Findeis D, Wiekenberg A, Mulcahy B, Luo L, Li Y, Lu Y et al. 2016. Neuroendocrine modulation sustains the C. elegans forward motor state. eLife 5.

Lorenzo R, Onizuka M, Defrance M, Laurent P. 2020. Combining single-cell RNA-sequencing with a molecular atlas unveils new markers for Caenorhabditis elegans neuron classes. Nucleic Acids Res.

Ma X, Zhao Z, Xiao L, Xu W, Kou Y, Zhang Y, Wu G, Wang Y, Du Z. 2021. A 4D single-cell protein atlas of transcription factors delineates spatiotemporal patterning during embryogenesis. Nature Methods 18: 893–902.

Masland RH. 2001. The fundamental plan of the retina. Nat Neurosci 4: 877–886.

Mazelova J, Ransom N, Astuto-Gribble L, Wilson MC, Deretic D. 2009. Syntaxin 3 and SNAP-25 pairing, regulated by omega-3 docosahexaenoic acid, controls the delivery of rhodopsin for the biogenesis of cilia-derived sensory organelles, the rod outer segments. J Cell Sci 122: 2003–2013.

Network BICC. 2021. A multimodal cell census and atlas of the mammalian primary motor cortex. Nature 598: 86–102.

Ojeda Naharros I, Gesemann M, Mateos JM, Barmettler G, Forbes A, Ziegler U, Neuhauss SCF, Bachmann-Gagescu R. 2017. Loss-of-function of the ciliopathy protein Cc2d2a disorganizes the vesicle fusion machinery at the periciliary membrane and indirectly affects Rab8-trafficking in zebrafish photoreceptors. PLoS Genet 13: e1007150.

Oren-Suissa M, Bayer EA, Hobert O. 2016. Sex-specific pruning of neuronal synapses in Caenorhabditis elegans. Nature 533: 206–211.

Poulin JF, Tasic B, Hjerling-Leffler J, Trimarchi JM, Awatramani R. 2016. Disentangling neural cell diversity using single-cell transcriptomics. Nat Neurosci 19: 1131–1141.

Reilly MB, Cros C, Varol E, Yemini E, Hobert O. 2020. Unique homeobox codes delineate all the neuron classes of C. elegans. Nature 584: 595–601.

Schindelin J, Rueden CT, Hiner MC, Eliceiri KW. 2015. The ImageJ ecosystem: An open platform for biomedical image analysis. Mol Reprod Dev 82: 518–529.

Schroeder NE, Androwski RJ, Rashid A, Lee H, Lee J, Barr MM. 2013. Dauer-specific dendrite arborization in C. elegans is regulated by KPC-1/Furin. Curr Biol 23: 1527–1535.

Serrano-Saiz E, Gulez B, Pereira L, Gendrel M, Kerk SY, Vidal B, Feng W, Wang C, Kratsios P, Rand JB et al. 2020. Modular Organization of Cis-regulatory Control Information of Neurotransmitter Pathway Genes in Caenorhabditis elegans. Genetics 215: 665–681.

Shaham S, Bargmann CI. 2002. Control of neuronal subtype identity by the C. elegans ARID protein CFI-1. Genes Dev 16: 972–983.

Shekhar K, Lapan SW, Whitney IE, Tran NM, Macosko EZ, Kowalczyk M, Adiconis X, Levin JZ, Nemesh J, Goldman M et al. 2016. Comprehensive Classification of Retinal Bipolar Neurons by Single-Cell Transcriptomics. Cell 166: 1308–1323 e1330.

Sun H, Hobert O. 2021. Temporal transitions in the post-mitotic nervous system of Caenorhabditis elegans. Nature 600: 93–99.

Taylor SR, Santpere G, Weinreb A, Barrett A, Reilly MB, Xu C, Varol E, Oikonomou P, Glenwinkel L, McWhirter R et al. 2021. Molecular topography of an entire nervous system. Cell 184: 4329–4347 e4323.

Vidal B, Aghayeva U, Sun H, Wang C, Glenwinkel L, Bayer EA, Hobert O. 2018. An atlas of Caenorhabditis elegans chemoreceptor expression. PLoS Biol 16: e2004218.

Vidal B, Santella A, Serrano-Saiz E, Bao Z, Chuang CF, Hobert O. 2015. C. elegans SoxB genes are dispensable for embryonic neurogenesis but required for terminal differentiation of specific neuron types. Development 142: 2464–2477.

Wang J, Kaletsky R, Silva M, Williams A, Haas LA, Androwski RJ, Landis JN, Patrick C, Rashid A, Santiago-Martinez D et al. 2015. Cell-Specific Transcriptional Profiling of Ciliated Sensory Neurons Reveals Regulators of Behavior and Extracellular Vesicle Biogenesis. Curr Biol 25: 3232–3238.

Wang J, Silva M, Haas LA, Morsci NS, Nguyen KC, Hall DH, Barr MM. 2014. C. elegans ciliated sensory neurons release extracellular vesicles that function in animal communication. Curr Biol 24: 519–525.

Ward S, Thomson N, White JG, Brenner S. 1975. Electron microscopical reconstruction of the anterior sensory anatomy of the nematode Caenorhabditis elegans. The Journal of comparative neurology 160: 313–337.

White JG, Southgate E, Thomson JN, Brenner S. 1986. The structure of the nervous system of the nematode Caenorhabditis elegans. Philosophical Transactions of the Royal Society of London B Biological Sciences 314: 1–340.

Yanowitz JL, Shakir MA, Hedgecock E, Hutter H, Fire AZ, Lundquist EA. 2004. UNC-39, the C. elegans homolog of the human myotonic dystrophy-associated homeodomain protein Six5, regulates cell motility and differentiation. Dev Biol 272: 389–402.

Yao Z, van Velthoven CTJ, Nguyen TN, Goldy J, Sedeno-Cortes AE, Baftizadeh F, Bertagnolli D, Casper T, Chiang M, Crichton K et al. 2021. A taxonomy of transcriptomic cell types across the isocortex and hippocampal formation. Cell 184: 3222–3241 e3226.

Yemini E, Lin A, Nejatbakhsh A, Varol E, Sun R, Mena GE, Samuel ADT, Paninski L, Venkatachalam V, Hobert O. 2021. NeuroPAL: A Multicolor Atlas for Whole-Brain Neuronal Identification in C. elegans. Cell 184: 272–288 e211.

Zeng H, Sanes JR. 2017. Neuronal cell-type classification: challenges, opportunities and the path forward. Nat Rev Neurosci 18: 530–546.

Zhang F, Bhattacharya A, Nelson JC, Abe N, Gordon P, Lloret-Fernandez C, Maicas M, Flames N, Mann RS, Colon-Ramos DA et al. 2014. The LIM and POU homeobox genes ttx-3 and unc-86 act as terminal selectors in distinct cholinergic and serotonergic neuron types. Development 141: 422–435.

