## Supplemental Information for "*C. elegans* Sine oculis/SIX-type homeobox genes act as homeotic switches to define neuronal subtype identities"

#### SUPPLEMENTAL FIGURE LEGENDS

**Suppl. Fig.S1: IL2 subtype markers predicted by scRNA analysis**
Single cell transcriptome data, displayed with the Cengen App (Taylor et al., 2021).
**A:** Single cell plot for each individual IL2 markers.
**B:** Expression of all members of the DEG/ENaC/ASIC family of ion channels throughout the *C. elegans* nervous system.

**Suppl. Fig.S2: NeuroPAL IDs and scoring for *unc-86* dependency of IL2 subtype markers**
**A:** NeuroPAL pictures for Fig 1F CRISPR reporters *flp-32(syb4374), aex-2(syb4447), dmsr-2(syb4514), flp-5(syb4513), npr-37(syb4440)*
**B:** Quantification of Fig 1 G *unc-86* dependent IL2 lateral markers *nlp-69(syb4512) egas-4 / otIs833* and IL2 dorsoventral markers *degl-2(syb5229) egas-1 / otIs846*, p-values by Fisher two-sided test.
**C:** *flp-32* IL2 DV and URA expression is dependent on *unc-86,* using the *flp-32(syb4374)* reporter in the *unc-86(ot1158)* null allele*. flp-32* is expressed at a much weaker level in IL2 L/R and was not scored here
**D:** *flp-5* IL2 DV expression is *unc-86* dependent, using the *flp-5(syb4513)* repoter in the *unc-86(ot1158)* null allele.

**Suppl. Fig.S3: *unc-39* autoregulation**
**A:** A 2.3kb *unc-39* promoter fusion (chromosomal integrant *otIs854)* displays diminished expression in both lateral IL2 and AIA in the *unc-39* mutants, with stronger effects in IL2 vs AIA, and in the *ok2137* null animals (which can only be scored as arrested L1) vs the *e257* hypomorph (which can be scored at later stages).
**B:** Molecular details of the *unc-39(syb4537$^{GFP}$o1193$^{bs_del}$)* putative binding site deletion.
**C:** An *unc-39(syb4537$^{GFP}$o1193$^{bs_del}$)* CRISPR homeodomain binding site deletion in the background our tagged *unc-39(syb4537)* allele affects adult expression in the lateral IL2 and, with a lower effect, in AIA. Late embryo / L1 worms express *unc-39* in a similar way.

**Suppl. Fig.S4: *unc-39* gene model and functional analysis.**
**A:** *unc-39* gene model with SIX and homeodomain protein domains, and mutant alleles (null *ok2137* and *unc-39*$^{R203Q}$ hypomorphs *e257, ot1171, ot1172* and *ot1173).*
**B:** Effect of *unc-39* on various IL2 fate markers *ins-1(syb5452) dmsr-2(syb4514) flp-32(syb4374)* (from **Fig.2E/F**), with NeuroPAL images included for cell identification purposes. Note that a pair of pharynx neurons (blue circle, with no NeuroPAL coloring) express *ins-1*.
**C:** Quantifications for **Fig.2E/F** marker losses with markers *nlp-69(syb4512[nlp-69::SL2::gfp::H2B]), ins-1(syb5452[ins-1::SL2::gfp::H2B]), flp-14(syb3323[flp-14::SL2::gfp::H2B]), degl-1 (otIs825)* and *egas-4 (otIs833)* for the lateral IL2; *dmsr-2(syb4514[dmsr-2::SL2::gfp::H2B]), flp-32(syb4374[flp-32::SL2::gfp::H2B]), degl-2(syb5229[degl-2::SL2::gfp::H2B])* and *egas-1(otIs846)* for the dorsoventral IL2*.* Dorsoventral marker gain was counted as all or nothing, reporting extra cell counts (0,1 or 2 cells past the usual 4 cells). For *flp-32* the very dim lateral IL2 wild type expression was not counted as proper expression, in the mutant IL2 are always brighter than this. Fisher test p-values are shown on top, number of worms on the bottom.
**D:** Pan-IL2 identity is not affected in the *unc-39* null or the *unc-39*$^{R203Q}$ hypomorphic allele, as assessed by unaffected expression of the *unc-17prom9* fragment (*otEx6986*), the *cho-1* fosmid reporter (*otIs354)*, the *klp-6* reporter *myIs13*. the IL1 *flp-3* reporter *otIs703*, the *unc-17*

reporter allele *syb4491* or the NeuroPAL color code (which also derives from *klp-6).*
**E:** Quantifications for **Fig.2E/F** *unc-39* misexpression experiments. Lateral markers, *egas-4 (otIs833)* and *nlp-69(syb4512[nlp-69::SL2::gfp::H2B]),* are fully off in *e257* control, but were counted as on even if only weakly expressed. Dorsoventral markers, *egas-1(otIs846) and degl-2(syb5229[degl-2::SL2::gfp::H2B])*, are counted as off if visibly weaker than in the *e257* control. Each dot is a worm, on the y-axis is the number of neurons expressing the marker. There are sometimes more neurons than just the six IL2 for *egas-4* and *nlp-69*. Wilcoxon rank sum tests were done for each line vs the hypomorph with p-values on top, number of worms on the bottom.
**F:** Effect of ectopic *unc-39* on *flp-32(syb4374[flp-32::SL2::gfp::H2B]),* expression (from **Fig.2E**), with NeuroPAL images included, as well as quantifications below for IL2 on the left, URA on the right. The *unc-39(ot1073)* animals are shown in a ventral view.

**Suppl. Fig.S5: *unc-39* affects IL2 morphology**
**A: Top:** same images as Fig.3A of a *klp-6 (myIs13)* cytoplasmic reporter. To better show fasciculation it is now depth-coded (orange is top of the stack, purple on bottom). On the left a wildtype image of the left/right pairs show up as yellow/purple, but on the right *unc-39* mutant panel, the right lateral IL2 joins the right ventral IL2 cell body and their dendrite fasciculate. **Bottom:** blow-up images.
**B:** GFP::CLA-1 punctae quantification, by taking the number of dendrites per worm that have multiple bright punctae before the cilia, like in the wild type lateral IL2. P-values are computed by Wilcoxon rank sum test.
**C:** Dauer arborization phenotype quantification. We scored if branching was similar to wild type and *e257* mutant animals, reduced or fully gone. We then scored whether we observed neurites projecting backward.
**D:** The same *srh-71* reporter *(otIs867)* is shown left to right at L3, in dauer and 24 hours after replating dauers on fresh food. Wildtype animals are on top, *unc-39* mutants at the bottom. Notice that after recovery the lateral IL2 are gone in the wild type, but are equally bright as the dorsoventral IL2.

**Suppl. Fig.S6: Quantification of *ceh-32* loss- and gain-of-function experiments**
**A:** Quantifications for the loss of *mgl-1* in the dorsoventral RMD across the two lines (*otEx7843*, *otEx7844*) and the gain of *lgc-37* in the lateral RMD (counting only extra cells beyond the usual two lateral RMD), in lines *otEx7688* and *otEx7689*. P-values were obtained by two-sided Fisher test.
**B:** Quantification for the misexpression experiments. Dorsoventral markers *nlp-45(ot1032)* and *unc-46 otIs568* are gained (only one lateral RMD at most for *unc-46).* The lateral markers *flp-19(syb3278)* and *nlp-11(syb4759)* are lost. Note that only one line is shown for *unc-46,* with only one lateral RMD being reprogrammed at most*;* another line *otEx7911* was considered but had no effect on the lateral RMD. P-values were obtained by Wilcoxon rank sum test.

**Suppl. Fig.S7: Characterization of the *C42D4.1/slrf-7* pan-RMD driver**
**A:** Genome structure of our *C42D4.1/slrf-7* promoter we use as a pan-RMD driver
**B:** Annotated close up of the *otEx7843* panel from **Fig.4C**, demonstrating that *slrf-7* is expressed in all RMDs. The animal is in a ventral position. The yellow triangles point to cells expressing both *slrf-7* and a *mgl-1* fragment known to be specific of the dorsoventral RMD subtype. The red triangles point to the lateral RMD, between the RMD of the previous subtype, with a brighter *slrf-7* expression in this subtype. White triangles point to the dorsal

SAAs, based on both the CeNGEN scRNA atlas and the fact that they make projections anteriorly in the sublaterals. The ventral SAA rarely express *slrf-7*, depending on transgenic lines.

**Suppl. Fig.S8: Effect of *unc-42* on *nlp-11* expression**
The lateral RMD subtype marker *nlp-11(syb4759)* becomes dim in the *unc-42(e419)* mutant. The lateral RMD are circled in red, neighbouring neurons are circled in blue. We used *unc-42* independent neurons like RMEV or RIH to assess brightness changes. Significance was calculated by Fisher two-sided test.

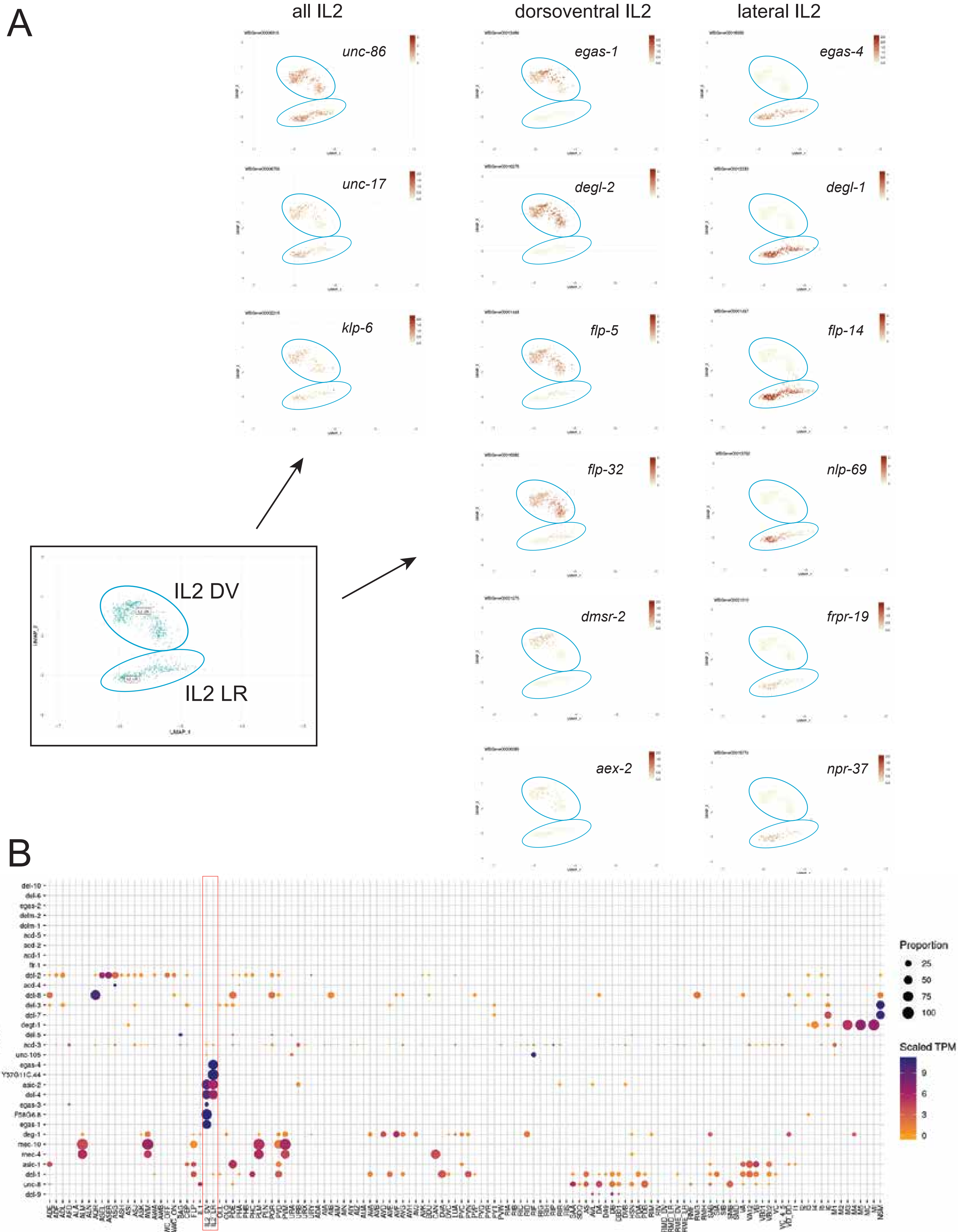
A
all IL2
dorsoventral IL2
lateral IL2
unc-86
unc-17
klp-6
egas-1
degl-2
flp-5
flp-32
dmsr-2
aex-2
egas-4
degl-1
flp-14
nlp-69
frpr-19
npr-37
IL2 DV
IL2 LR
B
Gene
Neuron
Proportion
25
50
75
100
Scaled TPM
9
6
3
0
del-10
del-6
egas-2
delm-2
delm-1
acd-5
acd-2
acd-1
flr-1
del-2
acd-4
del-8
del-3
del-7
degt-1
del-5
acd-3
unc-105
egas-4
Y57G11C.44
asic-2
del-4
egas-3
F58G6.8
egas-1
deg-1
mec-10
mec-4
asic-1
del-1
unc-8
del-9

Supp Fig. S1

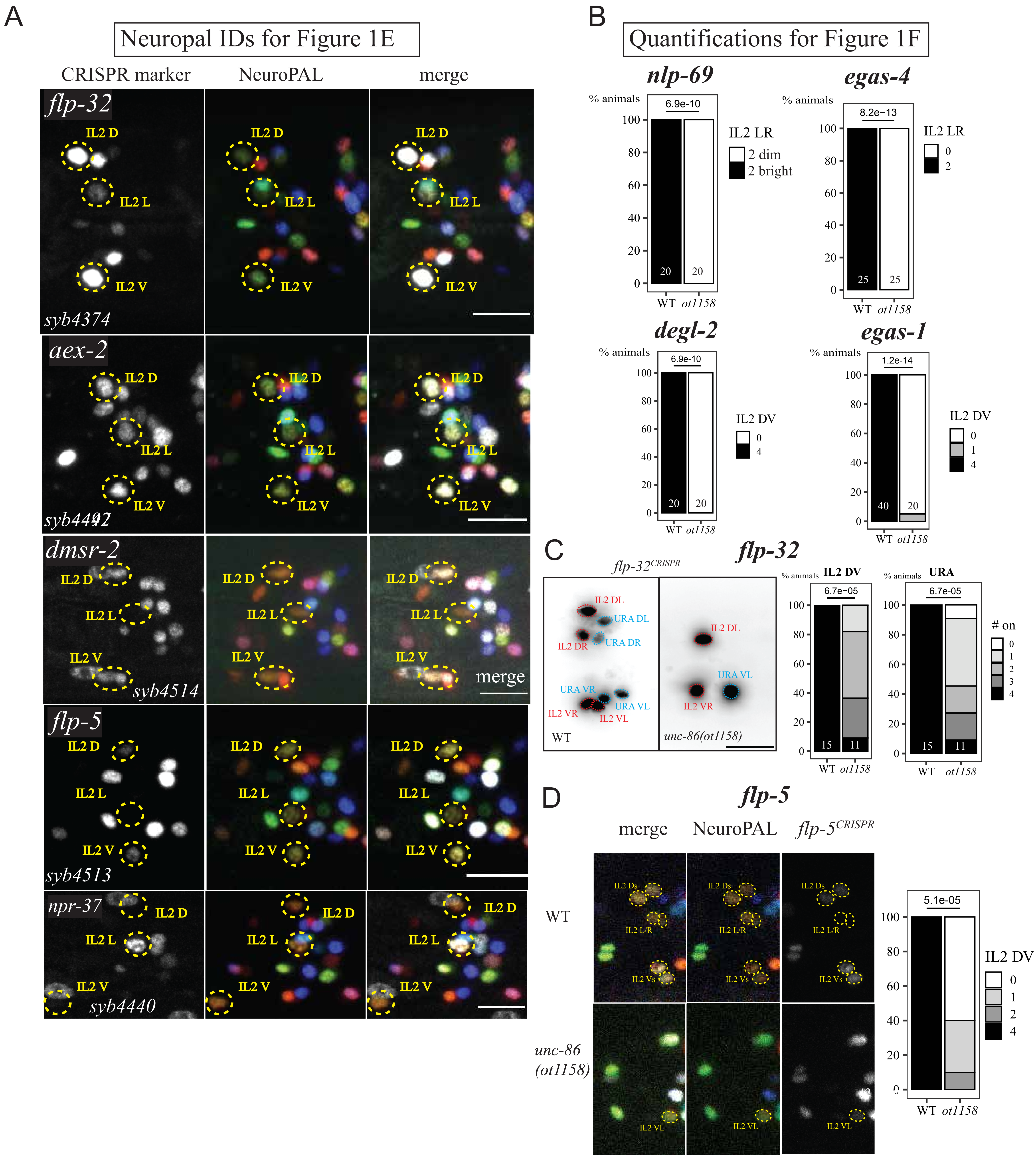

A
Neuropal IDs for Figure 1E
CRISPR marker
NeuroPAL
merge
flp-32
IL2 D
IL2 L
IL2 V
syb4374
aex-2
syb4492
dmsr-2
syb4514
flp-5
syb4513
npr-37
syb4440
B
Quantifications for Figure 1F
nlp-69
% animals
6.9e-10
IL2 LR
2 dim
2 bright
WT
ot1158
egas-4
8.2e-13
degl-2
IL2 DV
egas-1
1.2e-14
C
flp-32
flp-32CRISPR
IL2 DL
URA DL
IL2 DR
URA DR
URA VR
URA VL
IL2 VR
IL2 VL
unc-86(ot1158)
6.7e-05
URA
# on
D
flp-5
merge
NeuroPAL
flp-5CRISPR
WT
unc-86
(ot1158)
IL2 Ds
IL2 L/R
IL2 Vs
5.1e-05


Supp Fig. S2

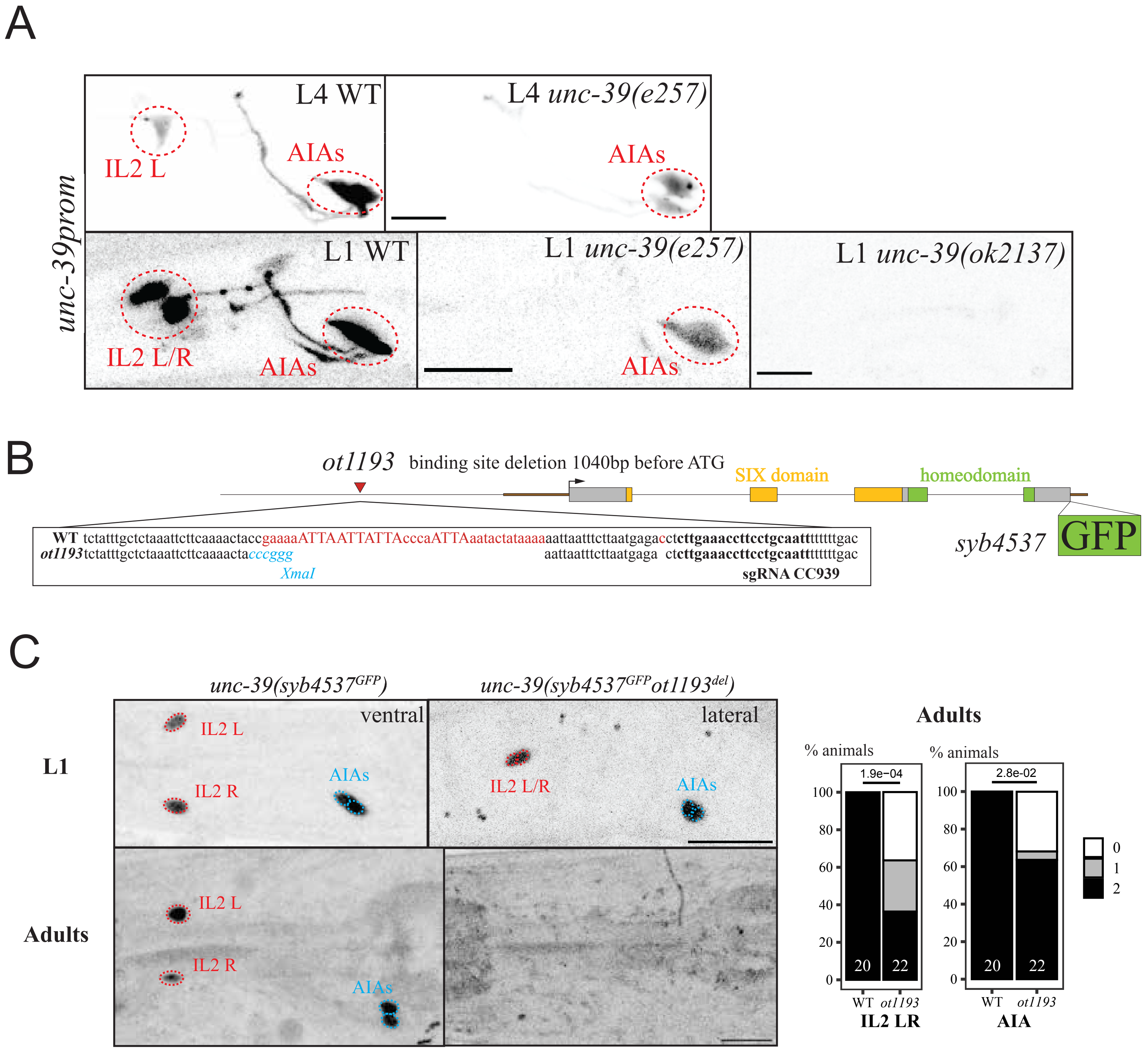
A
unc-39prom
L4 WT
IL2 L
AIAs
L4 unc-39(e257)
AIAs
L1 WT
IL2 L/R
AIAs
L1 unc-39(e257)
AIAs
L1 unc-39(ok2137)
B
ot1193 binding site deletion 1040bp before ATG
SIX domain
homeodomain
WT tctatttgctctaaattcttcaaaactaccgaaaaATTAATTATTAcccaATTAaatactataaaaaattaatttcttaatgagacctcttgaaaccttcctgcaattttttttgac
ot1193 tctatttgctctaaattcttcaaaactacccgggg aattaatttcttaatgaga ctcttgaaaccttcctgcaattttttttgac
XmaI
sgRNA CC939
syb4537
GFP
C
unc-39(syb4537GFP)
unc-39(syb4537GFPot1193del)
ventral
lateral
L1
IL2 L
IL2 R
AIAs
IL2 L/R
AIAs
Adults
IL2 L
IL2 R
AIAs
Adults
% animals
% animals
1.9e−04
2.8e−02
100
80
60
40
20
0
20
22
20
22
0
1
2
WT
ot1193
IL2 LR
WT
ot1193
AIA

Supp Fig. S3

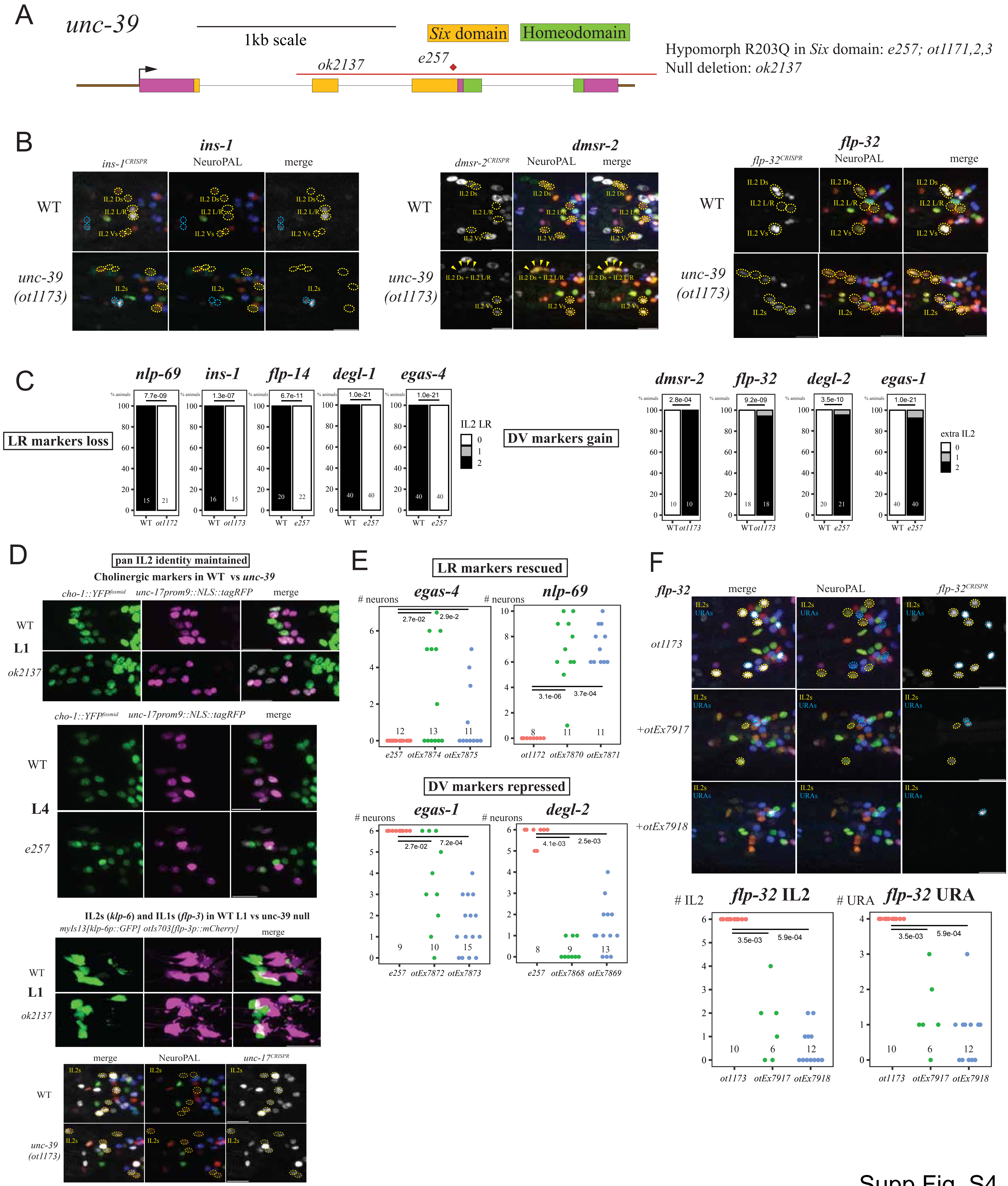
A
unc-39
1kb scale
Six domain
Homeodomain
ok2137
e257
Hypomorph R203Q in Six domain: e257; ot1171,2,3
Null deletion: ok2137
B
ins-1
ins-1CRISPR
NeuroPAL
merge
WT
unc-39 (ot1173)
IL2 Ds
IL2 L/R
IL2 Vs
IL2s
dmsr-2
dmsr-2CRISPR
IL2 Ds + IL2 L/R
flp-32
flp-32CRISPR
C
LR markers loss
nlp-69
ins-1
flp-14
degl-1
egas-4
% animals
7.7e-09
1.3e-07
6.7e-11
1.0e-21
1.0e-21
WT
ot1172
ot1173
e257
IL2 LR
DV markers gain
dmsr-2
flp-32
degl-2
egas-1
2.8e-04
9.2e-09
3.5e-10
1.0e-21
extra IL2
D
pan IL2 identity maintained
Cholinergic markers in WT vs unc-39
cho-1::YFPfosmid
unc-17prom9::NLS::tagRFP
merge
L1
ok2137
L4
e257
IL2s (klp-6) and IL1s (flp-3) in WT L1 vs unc-39 null
myIs13[klp-6p::GFP] otIs703[flp-3p::mCherry]
unc-17CRISPR
E
LR markers rescued
egas-4
nlp-69
### neurons
2.7e-02
2.9e-2
3.1e-06
3.7e-04
otEx7874
otEx7875
otEx7870
otEx7871
DV markers repressed
egas-1
degl-2
7.2e-04
4.1e-03
2.5e-03
otEx7872
otEx7873
otEx7868
otEx7869
F
+otEx7917
+otEx7918
URAs
# IL2
flp-32 IL2
### URA
flp-32 URA
3.5e-03
5.9e-04

Supp Fig. S4

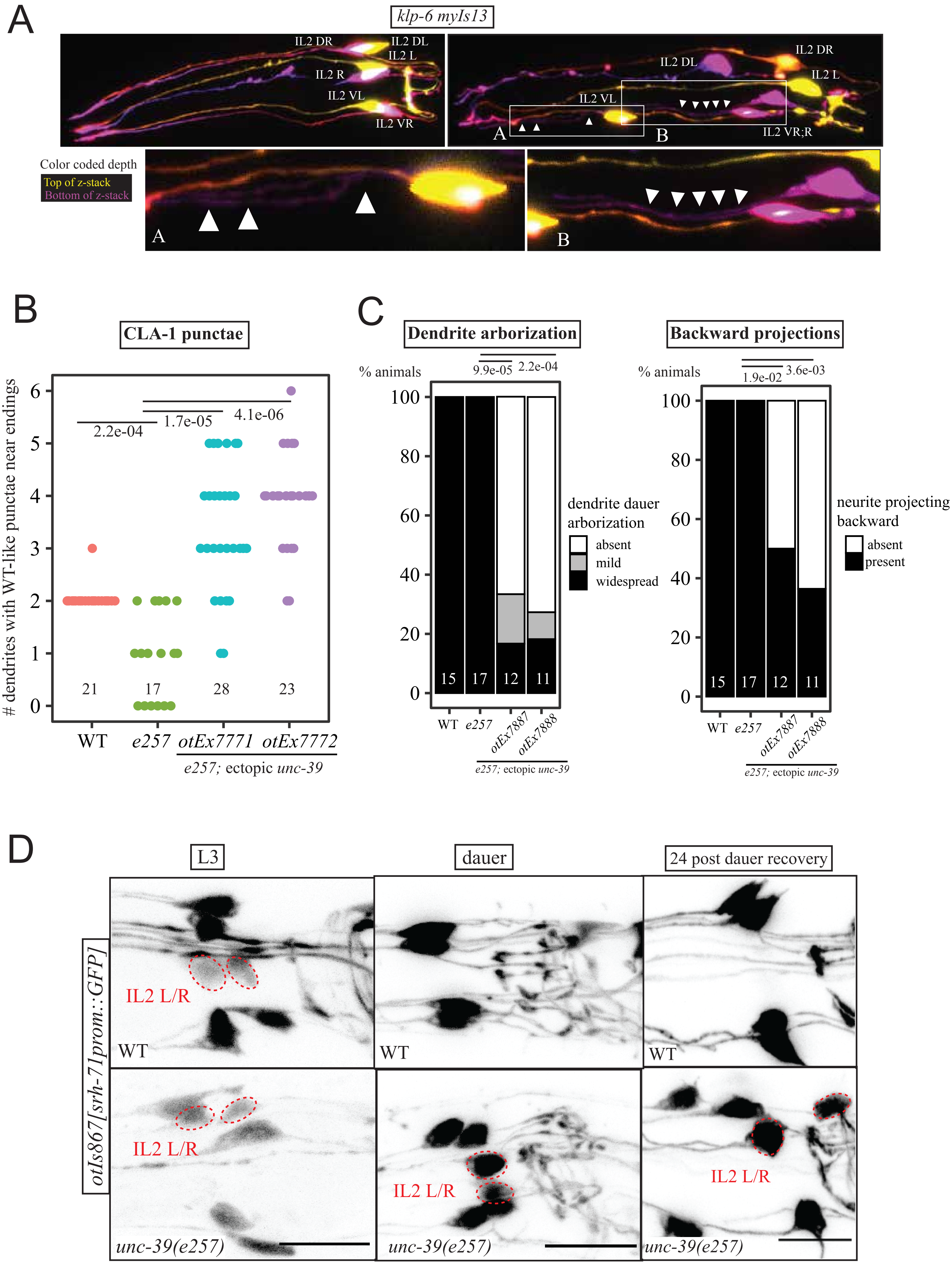
A
klp-6 myIs13
IL2 DR
IL2 DL
IL2 L
IL2 R
IL2 VL
IL2 VR
IL2 DL
IL2 DR
IL2 L
IL2 VL
IL2 VR;R
Color coded depth
Top of z-stack
Bottom of z-stack
B
CLA-1 punctae
### dendrites with WT-like punctae near endings
2.2e-04
1.7e-05
4.1e-06
21
17
28
23
WT
e257
otEx7771
otEx7772
e257; ectopic unc-39
C
Dendrite arborization
% animals
9.9e-05
2.2e-04
dendrite dauer arborization
absent
mild
widespread
15
17
12
11
WT
e257
otEx7887
otEx7888
e257; ectopic unc-39
Backward projections
% animals
1.9e-02
3.6e-03
neurite projecting backward
absent
present
D
L3
dauer
24 post dauer recovery
otIs867[srh-71prom::GFP]
IL2 L/R
WT
unc-39(e257)

Supp Fig. S5

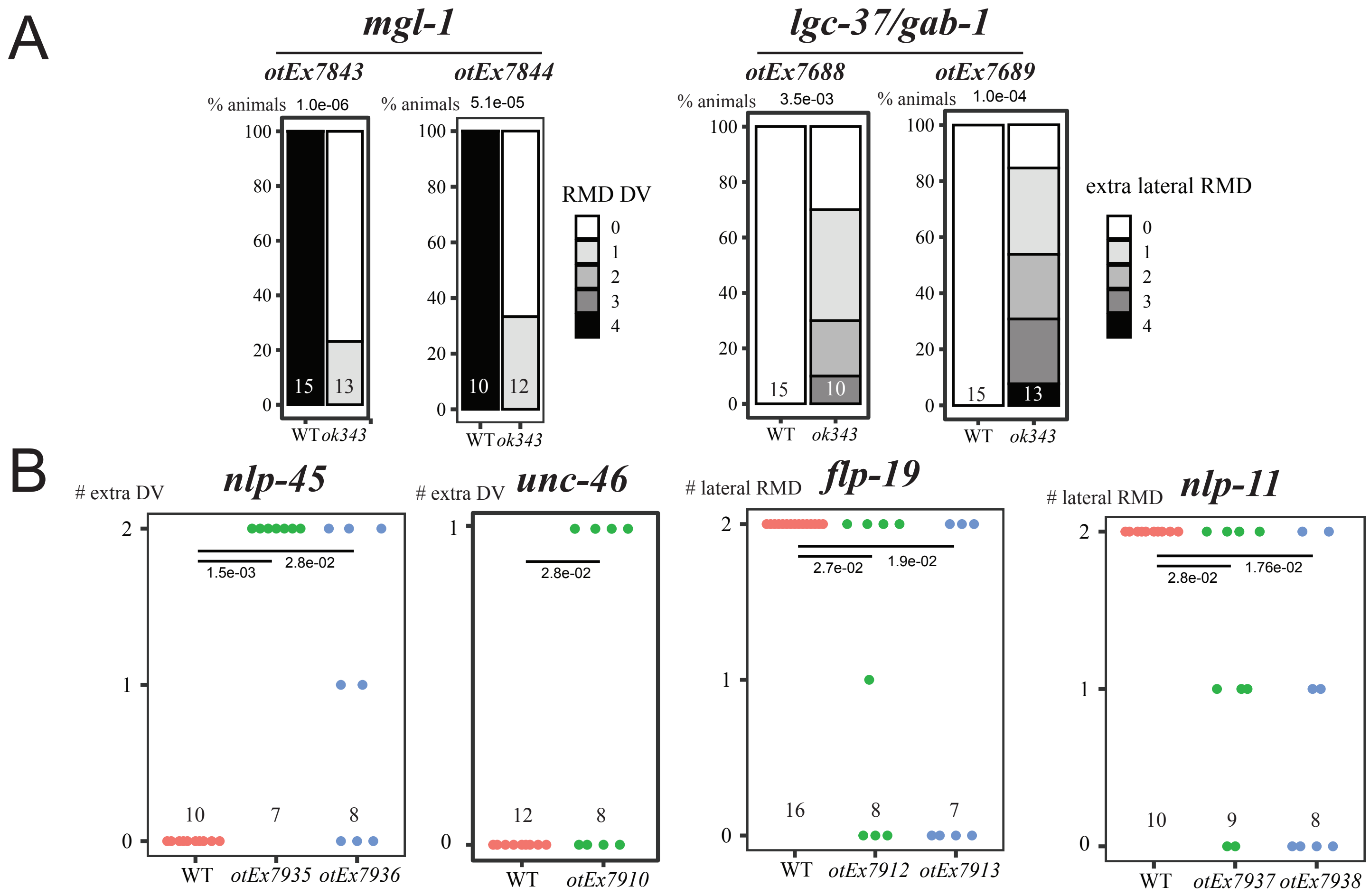
A
mgl-1
otEx7843
otEx7844
% animals 1.0e-06
% animals 5.1e-05
RMD DV
lgc-37/gab-1
otEx7688
otEx7689
% animals 3.5e-03
% animals 1.0e-04
extra lateral RMD
WT ok343
B
### extra DV nlp-45
1.5e-03
2.8e-02
WT otEx7935 otEx7936
### extra DV unc-46
2.8e-02
WT otEx7910
### lateral RMD flp-19
2.7e-02
1.9e-02
WT otEx7912 otEx7913
### lateral RMD nlp-11
2.8e-02
1.76e-02
WT otEx7937 otEx7938

Supp Fig. S6

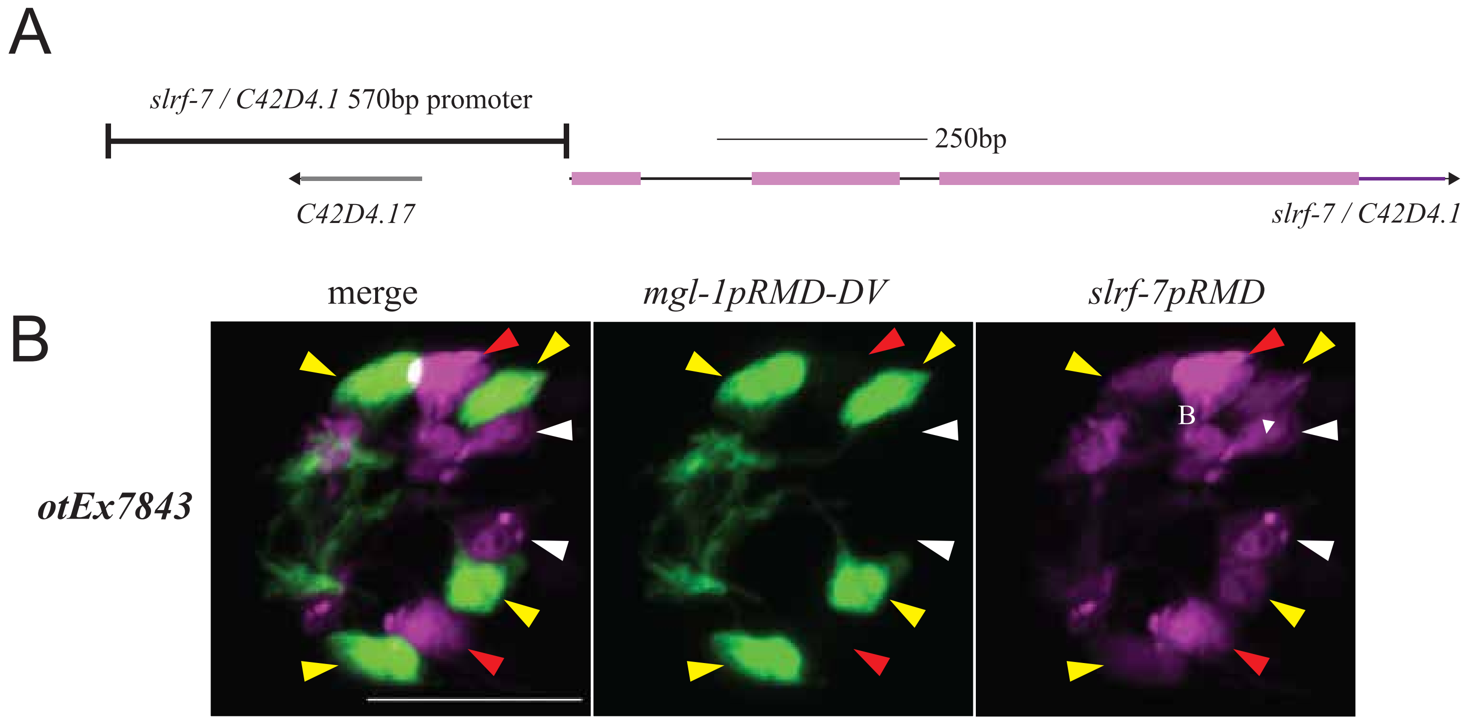
A
slrf-7 / C42D4.1 570bp promoter
250bp
C42D4.17
slrf-7 / C42D4.1
B
merge
mgl-1pRMD-DV
slrf-7pRMD
otEx7843
B

Supp Fig. S7

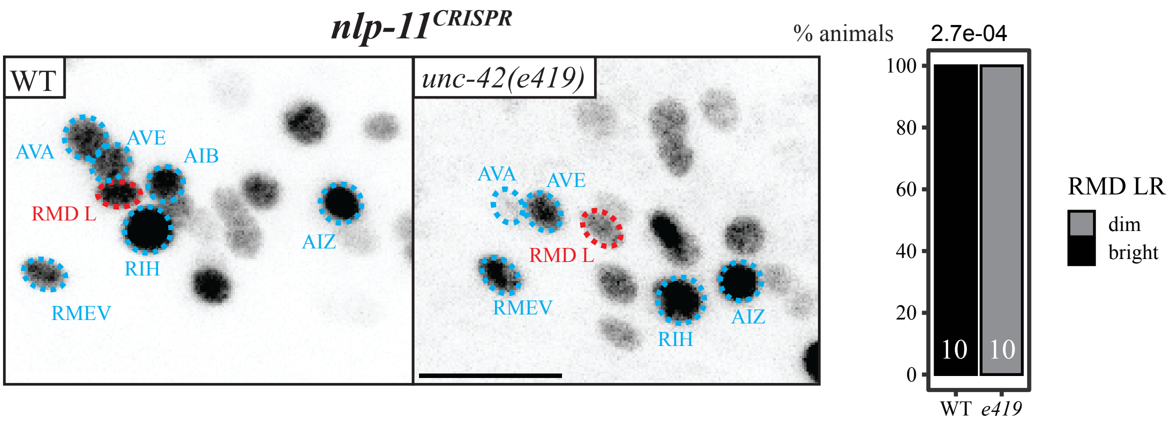
nlp-11CRISPR
WT
unc-42(e419)
AVA
AVE
AIB
RMD L
RIH
AIZ
RMEV
AVA
AVE
RMD L
RMEV
RIH
AIZ
% animals
2.7e-04
100
80
60
40
20
0
10
10
WT
e419
RMD LR
dim
bright

Supp Fig. S8
